# Visual objects refine head direction coding

**DOI:** 10.1101/2024.10.21.619417

**Authors:** Dominique Siegenthaler, Henry Denny, Sofía Skromne Carrasco, Johanna Luise Mayer, Daniel Levenstein, Adrien Peyrache, Stuart Trenholm, Émilie Macé

## Abstract

Animals use visual objects to guide navigation-related behaviors, from hunting prey, to escaping predators, to exploring the world. However, little is known about where visual objects are encoded in the mouse brain or how they impact processing in the spatial navigation system. Using functional ultrasound imaging in mice, we conducted a brain-wide screen and identified brain areas that were preferentially activated by images of objects compared to scrambled versions of the same images. While visual cortical areas did not show a significant preference, regions associated with spatial navigation were preferentially activated by visual objects. Electrophysiological recordings in postsubiculum, the primary cortical area of the head direction (HD) system, further confirmed a preference for visual objects, which was present in both HD cells and fast-spiking interneurons. Finally, we found that visual objects dynamically modulated HD cells, selectively increasing firing rates of HD cells aligned with a visual landmark’s direction, while decreasing activity in HD cells coding for other directions. These results reveal that visual objects refine population-level coding of head direction.

## Introduction

The spatial navigation system is comprised of neurons whose firing rate is modulated by variables including the direction an animal is facing and the location it occupies in an environment^1–3^. In turn, these neurons are believed to support the cognitive map^4,5^. However, in addition to spatial variables, neurons within the spatial navigation system are modulated by visual inputs^6^, as visual objects^7^ can serve as environmental landmarks^8^. Moving a visual landmark in an environment can result in a corresponding shift in the tuning of head direction (HD) cells^9^, place cells^10,11^, and grid cells^3^. Furthermore, placing an animal in the dark can lead to tuning instability^12,13^. Thus, visual landmarks can anchor the internal representation of space^14^. However, other than helping orientate the tuning of these cells, much remains to be known about how visual objects modulate the firing properties of neurons within the spatial navigation system.

A detailed understanding of how the brain encodes visual objects has been gleaned from decades of work, primarily in humans and non-human primates^15^. Briefly, spanning from primary visual cortex (V1) to inferotemporal cortex, there are a series of brain areas arranged in a hierarchical manner, referred to as the ‘ventral visual stream’, tasked with encoding increasingly high-level representations of visual objects^15,16^. Along these lines, studies in rodents have found that lateral visual cortical areas appear to exhibit certain ventral-steam-like properties^17–20^ and that rodents are able to adapt their behaviors in response to visual objects^18,21,22^. However, a clear understanding of how and where visual objects are encoded in the rodent brain is largely missing.

To address these issues, we directly examined where representations of visual objects can be detected in the mouse brain and how visual objects impact processing within the spatial navigation system. First, we performed a brain-wide screen, using functional ultrasound (fUS) imaging^23–25^, to look for areas activated more strongly by visual objects than by scrambled versions of the same images. While we did not find a preference for visual objects in the mouse’s ventral visual cortical areas, we found an enrichment of visual object preferring areas within the spatial navigation system, which we then validated with single-cell electrophysiology. Next, focusing on one of the top hits of our fUS screen, the postsubiculum (PoSub), the cortical hub of the HD system, we examined the firing properties of neurons in freely moving mice and found that a preference for visual objects was present in both HD cells and fast spiking (FS) interneurons. Finally, we found that visual objects act to dynamically refine population coding of HD in PoSub, increasing the firing rate of HD cells whose preferred firing directions correspond to a visual landmark, while decreasing the firing rate of HD cells coding other directions.

## Results

### A brain-wide screen for areas that preferentially respond to visual objects

We set out to screen for brain areas in the mouse that were preferentially modulated by visual objects. Functional brain imaging experiments in humans, comparing activity driven by the presentation of visual objects versus scrambled versions of the same images, were important in revealing the importance of ventral stream cortical areas in encoding visual objects^26,27^. Inspired by those experiments, here we performed a brain-wide screen in mice using fUS imaging^23–25^ to search for areas that responded preferentially to visual objects. fUS is a neuroimaging method that monitors changes in blood volume, a proxy for neuronal activity^28^, and enables brain-wide volumetric recordings with a spatial resolution of ∼250 μm and a temporal resolution of 2 Hz^29^.

Mice were implanted with a COMBO cranial window^30^ and head-fixed under the ultrasound probe (**Fig. 1a)**. Visual stimuli were presented while fUS signals were measured simultaneously from almost the whole mouse brain (excluding olfactory bulbs, cerebellum and parts of the hindbrain). Images were presented 60° in height, centered along the horizontal meridian and 20° above the vertical meridian (**Fig. 1a**), corresponding to the ‘focea’, a part of the visual field thought to be particularly behaviorally relevant for mice^31^ and that has been hypothesized to be important for processing visual landmarks^32^. Images of 48 different objects were shown, covering a range of object categories (**Extended Data Fig. 1**). Each image contained a single object centered on a grey background and was shown for 0.5s. For image scrambling, we used a texture scrambling method^33^ (which we refer to as Scrambled_T_) that has previously been used in studies of the primate ventral visual stream^34,35^. The 48 images were split into 4 blocks of 12 images, and 4 blocks of corresponding scrambled images (**Fig. 1b, Extended Data Fig. 1**).

**Fig 1.**
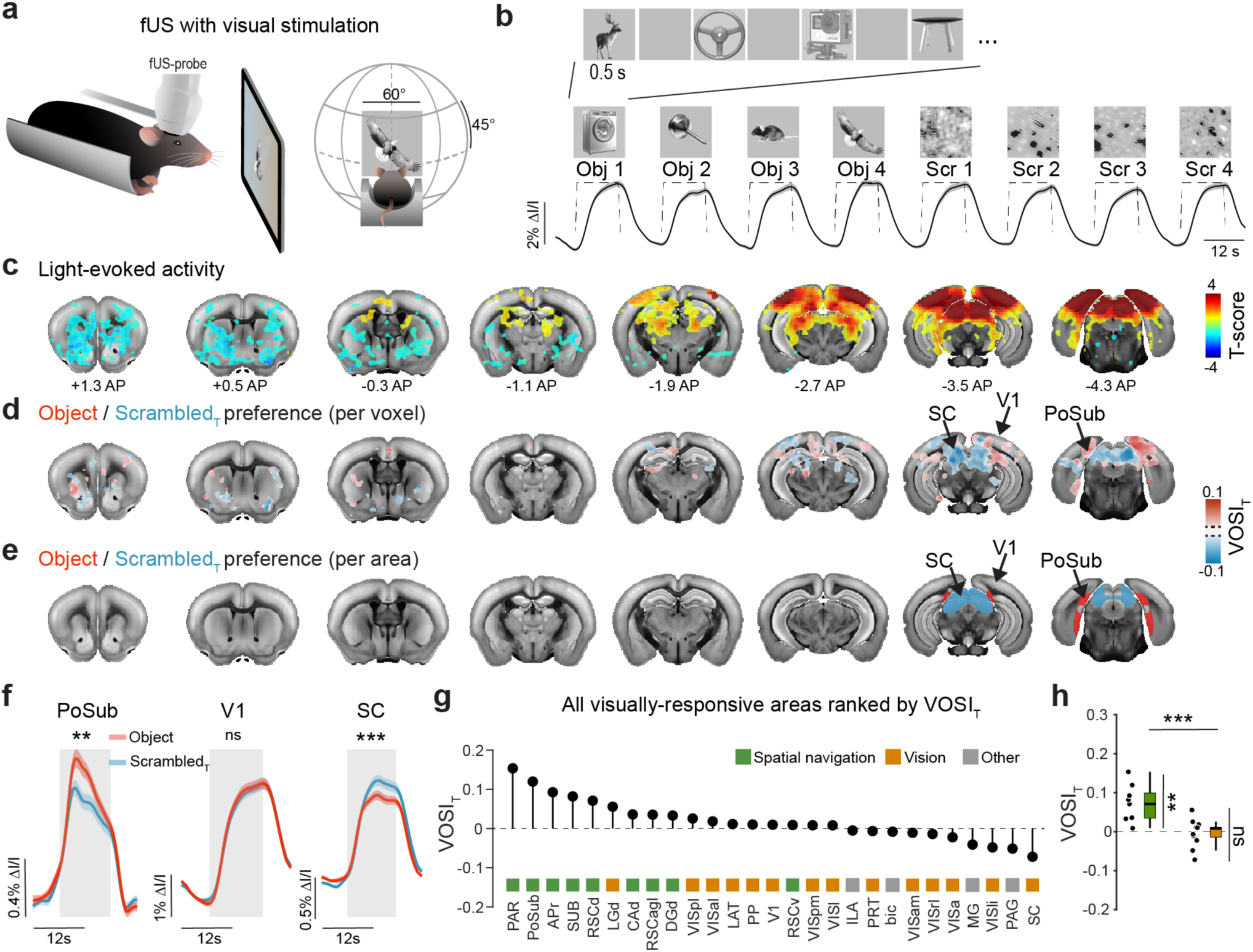
Visual object sensitive brain areas revealed with brain-wide fUS imaging. **a**, Schematic of fUS imaging with visual stimulation. **b**, *top*, Schematic of the stimulus design, with blocks of object and scrambled images, interleaved with full-field grey stimuli. **b,** *bottom*, Pooled fUS responses (mean ± s.e.m.) from V1 for each stimulus block (n = 56 sessions from 7 anesthetized animals; same data for all analyses in this figure). **c**, Visually-evoked fUS responses (all image blocks as regressors) overlaid on coronal brain images at indicated positions. Only T-scores significantly different from zero are shown, p < 0.05, FDR-corrected, mixed effects model. **d**, Visually-active voxels color coded by VOSIT values are displayed on coronal brain images (only absolute VOSIT values larger than 0.015 are shown). **e**, Brain areas with a VOSIT significantly different from zero are colored by their VOSIT values, p < 0.05, FDR-corrected, mixed effects model. **f**, Average fUS responses to Object and ScrambledT images, from PoSub, V1 and SC. p (from left to right) = 0.001, 0.359, 6.3*10^-7^, Bonferroni-Holm (B-H) corrected, mixed effects model. **g**, Rank ordering of visually-responsive brain areas according to VOSIT values (*black dots*), color-coded (*squares*) according to Spatial navigation (*green*), Vision (*orange*) or Other (*grey*) brain networks. **h**, VOSIT values of all brain regions belonging to either Spatial navigation (*green*) or Vision networks (*orange*). For statistical comparison between the two networks, p =7 .5*10^-4^, Mann-Whitney U-test. VOSIT values for Spatial navigation but not Vision networks statistically differed from 0. p(Spatial navigation) = 0.008, p(Vision) = 0.952, B-H corrected, one-sample Wilcoxon signed rank test.

First, we examined the visual response, in anesthetized mice, to all 8 blocks (i.e. combining object and scrambled stimuli). We found visual stimulus evoked activity in voxels throughout the mouse visual system, including visual cortex, superior colliculus, and lateral geniculate nucleus of the thalamus (**Fig. 1c**). After anatomically segmenting the brain into ∼100 areas, we found visual activity in 27 brain areas (**Extended Data Fig. 1**). Next, we computed a visual object sensitivity index (VOSI_T_), to quantify preferences for object vs. texture scrambled images. Positive VOSI_T_ values indicate stronger responses for object stimuli and negative VOSI_T_ indicates stronger responses for scrambled stimuli. We noted an enrichment in positive VOSI_T_ voxels in the hippocampal formation and retrosplenial cortex (RSC; **Fig. 1d, Extended Data Fig. 1**). VOSI_T_ analysis at the scale of brain areas revealed a small number of brain areas with a statistically significant preference for visual objects: the PoSub, the parasubiculum, and the area prostriata (**Fig. 1d-f**). In contrast, many areas, including V1 did not exhibit a preference for objects or scrambled images (**Fig. 1d-f**). We found one area, the superior colliculus (SC), responded significantly more strongly to scrambled images (**Fig. 1d-f**). Importantly, we found similar results when we repeated experiments in awake animals (**Extended Data Fig. 2**).

Next, to examine if there was a systematic bias for visual object sensitivity at the brain network-level, we rank-ordered visually responsive brain areas according to their VOSI_T_ values. Then, based on literature review, we categorized brain areas as belonging to either 1) the spatial navigation system, 2) the visual system or 3) other (**Fig. 1g**, see **Supplementary Table 1** for the list of brain areas and their assigned systems). Within the top ten areas with highest VOSI_T_ values, eight are closely associated with the spatial navigation system (**Fig. 1g**). Consistent with this, the average VOSI_T_ value of all spatial navigation related areas was significantly larger than zero (**Fig. 1h**). We found a similar result—that a preference for visual objects over scrambled images was restricted to spatial navigation areas—when we clustered brain areas, or visually responsive pixels, based solely on their object and scrambled response kernels (**Extended Data Fig. 3**).

Finally, as some studies indicate that, similar to primates, visual object processing in mouse cortex is biased towards ventral visual cortical areas^17,18^, we subdivided the ‘visual system’ group into ventral and dorsal visual cortical areas. However, we did not detect a difference in activation by object versus scrambled images for either ventral or dorsal visual cortical areas (**Extended Data Fig. 4**). Taken together, these results reveal a preference for visual objects within the mouse’s spatial navigation system, not its visual system.

### A preference for visual objects in postsubicular neurons

To ensure that visual object preferences identified with fUS were driven by neuronal firing^28^, we performed acute electrophysiological recordings from several brain areas in head-fixed anesthetized mice (**Fig. 2a, Extended Data Fig. 5**). We recorded from regions of the spatial navigation system: PoSub (which had a positive VOSI_T_ in fUS), as well as dorsal and ventral retrosplenial cortex (RSCd and RSCv). We chose RSC because from our fUS experiments RSCd, but not RSCv, trended towards preferring object images, and RSCd was among the most object preferring regions outside the subiculum. Additionally, previous work suggests RSC is a spatial navigation area involved with processing visual landmarks^36,37^. As control regions, we recorded from two visual areas: V1 (which did not exhibit a preference in fUS experiments), and SC (which had a texture preference in fUS experiments). We presented animals with the same object images as in fUS experiments, but now included a second form of scrambling. The texture scrambling method used above corrupts object identity but can also introduce local contrast changes into background areas that were previously solid gray. To control for this, taking advantage of the larger number of trials possible with electrophysiology, we added a second scrambling method, diffeomorphic transformation^38^, which maintains overall object shape while modifying local pixel- level spatial organization (Scrambled_D_; **Fig. 2b; Extended Data Fig. 6**).

**Fig 2.**
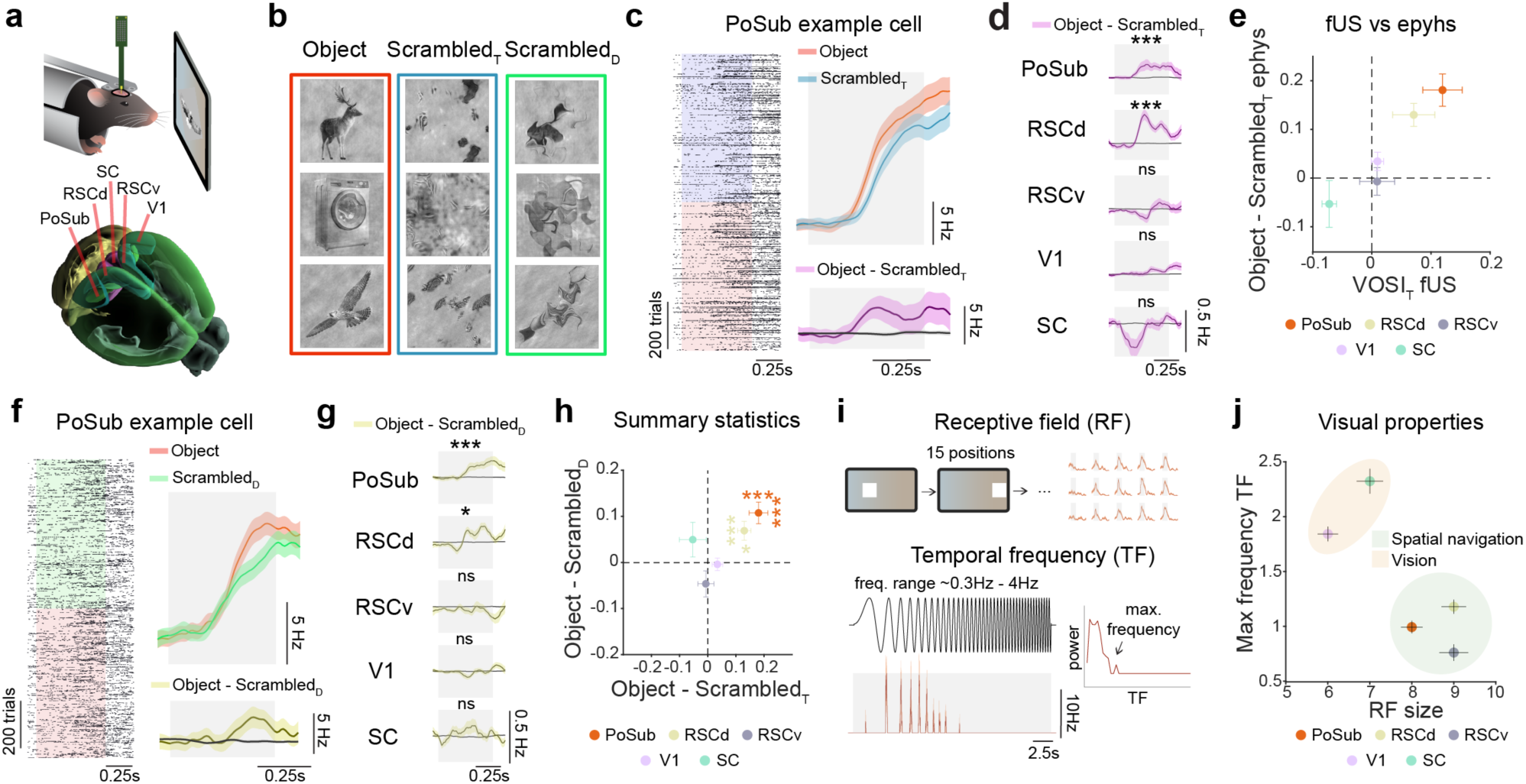
A preference for visual objects in postsubiculum and dorsal retrosplenial cortex in single cell recordings. **a**, Schematic of the experimental approach and the recording sites **b**, *left*, Example Object (*red box*), ScrambledT (texture; *blue box*) and ScrambledD (diffeomorphic transformations; *green box*) images. **c**, *left*, Raster plot showing spiking responses from a single neuron in PoSub to Object (*red*) and ScrambledT (*blue*) images. **c**, *right top*, Mean (*solid line*) *±* s.e.m. (*shaded area*) responses to Object (*red*) and ScrambledT (*blue*) images from the same cell. **c**, *right bottom*, Mean (*solid line*) *±* s.e.m. (*shaded area*) response difference of Object-ScrambledT responses. Black solid line with shaded area shows the mean *±* s.e.m. for 100 permutations. **d**, Traces of Object-ScrambledT (*purple line,* mean ± s.e.m. across cells). A shuffled response distribution is shown for visualization purposes (*black line,* mean ± s.e.m. of 100 permutations of trial labels shuffling), p (from top to bottom) = 2.4*10^-8^, 2.4*10^-5^, 0.595, 0.511, 0.351, B-H corrected, Wilcoxon signed rank test, (n (from top to bottom) = 192, 120, 81, 145, 61 cells). **e**, Object preference compared to texture stimuli for all five regions calculated using fUS data (VOSIT, x-axis, mean *±* s.e.m. n = 57 sessions) and ephys data (Object-ScrambledT, y-axis, mean *±* s.e.m., n’s as in (d)). **f-g**, The same as (**c-d**), except for Object vs. ScrambledD, p (from top to bottom) = 2.2*10^-6^, 0.017, 0.595, 0.456, 0.456, B-H corrected, Wilcoxon signed rank test, (n’s as in d). **h**, Object-ScrambledT and Object-ScrambledD values for neurons from the five different brain areas (mean *±* s.e.m., n’s and p’s as in (d,g)). **i**, Schematic of the visual stimulus protocols for measuring receptive field size, and spatial and temporal frequency preferences. **j**, Results of the receptive field and temporal frequency experiments described in (i) for all five regions (mean *±* s.e.m., see **Extended data Fig. 7** for details).

First, we compared fUS recordings to single unit spike recordings, focusing on the difference between responses to objects and texture scrambled images (**Fig. 2c-e**). For single cell responses, we tested for a preference for visual objects by subtracting the response to Scrambled_T_ images from the response to Objects (Object-Scrambled_T_). We found that neurons in PoSub and RSCd exhibited a preference for visual objects. Neurons in SC, V1 and RSCv cortex did not exhibit a preference for visual objects, though in accordance with the fUS results there was a negative trend in the SC Object-Scrambled_T_ response (**Fig. 2d**). For all five regions, spiking recapitulated fUS preferences for object versus texture scrambled images, with a strong correlation between the object preferences calculated with both recording modalities (**Fig. 2e**, R = 0.97). Next, we examined responses to diffeomorphic transformations (**Fig. 2f**). From this point forward, we considered brain areas to prefer visual objects if they showed a preference for Objects over both Scrambled_T_ and Scrambled_D_. This was the case for PoSub and RSCd (**Fig. 2f-h**). In contrast, we found that V1 and RSCv cortex did not show a preference for objects (**Fig. 2g,h**). Lastly, SC neurons did not show a preference for diffeomorphic scrambles, suggesting that the preference for texture scrambled images in fUS recordings is more related to the texture scrambling method itself than to the disruption of object identity (**Fig. 2g,h**).

Next, we wondered if PoSub and RSCd exhibited other visual feature preferences that differentiated them from non-visual object preferring areas. We measured additional visual properties (receptive field size, spatial and temporal frequency tuning) of neurons from the five brain areas outlined above. We found that neurons in PoSub and RSCd exhibited larger receptive fields and lower temporal frequency preferences compared to SC and V1, but had similar tuning properties to non-object preferring neurons in RSCv (**Fig 2i,j**; **Extended Data Fig. 7**).

### In PoSub, head-direction and fast spiking cells exhibit a preference for visual objects

How do visual objects modulate the processing of spatial information? As PoSub was one of the most visual object preferring regions in our screen, and as it is known to possess HD cells^2,39^ and be modulated by visual landmarks^14^, we focused further on this region. To obtain freely-moving recordings and head-fixed visual stimulation recordings from the same neurons, we chronically implanted a silicon probe in PoSub (**Extended Data Fig. 8**). We then introduced mice to an open field arena, with one visual landmark on the wall, for freely moving recording sessions (**Fig. 3a**). Immediately following open field recordings, animals were kept awake, head-restrained, and shown the same set of visual stimuli as outlined above for anesthetized animals (**Fig. 3a**).

**Fig 3.**
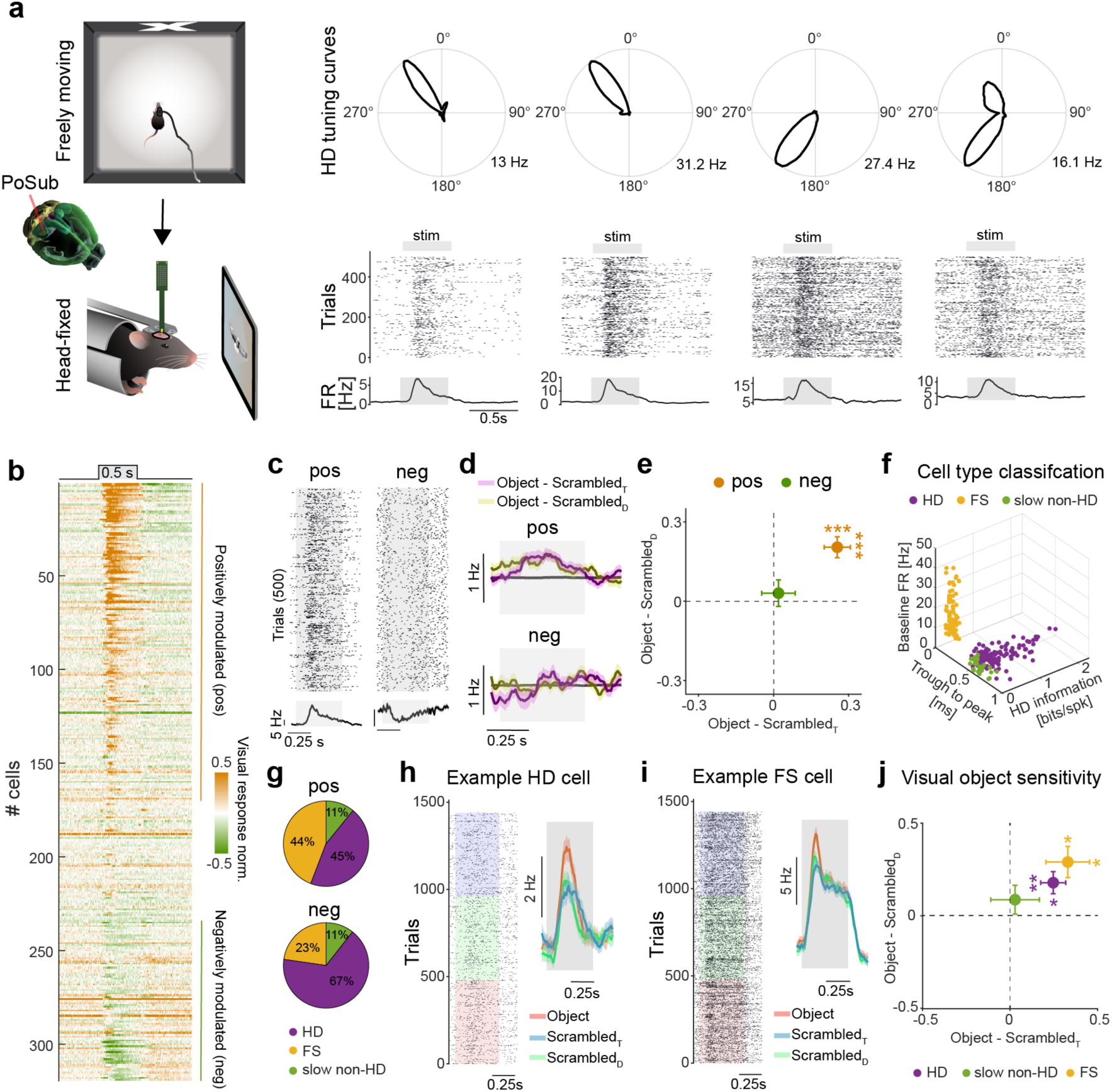
A preference for visual objects is present in HD and FS neurons in PoSub. **a**, *left*, Schematic of the two awake experimental paradigms, involving freely moving exploration of an open-field arena (*top*) followed by head-fixation and presentation of visual stimuli (*bottom*). **a**, *right,* Polar plots for four recorded HD cells in PoSub (*top*), and responses of the same cells to 500 randomly selected visual stimulus presentations (*bottom*), plotted both as raster plots and as a mean visually-evoked firing rate (FR). **b**, The average visually-evoked responses of all cells recorded in PoSub in awake animals (n = 319 cells from 5 animals), ordered from top-to-bottom by visual response strength. **c**, Example raster plots and average firing rates of an example neuron that was positively modulated by visual stimulation (*left*) and an example neuron that was negatively modulated by visual stimulation (*right*). Shown are 500 randomly selected trials on the raster plot and mean firing rate across all trials below. **d**, Mean (*solid line*) ± s.e.m. (*shaded area*) across positively or negatively modulated cells for Object-ScrambledT and Object-ScrambledD responses (n (from top to bottom) = 170, 86 cells). Black solid line with shaded area shows the mean ± s.e.m. for 100 permutations. **e**, Object-ScrambledT and Object-ScrambledD values for positively and negatively visually modulated PoSub neurons (mean ± s.e.m.), p (VOSIT, from left to right) = 8.56*10^-6^, 0.98, p (VOSID, from left to right) = 2.2*10^-5^, 0.9, B-H corrected, Wilcoxon signed rank test, (n (from top to bottom) = 170, 86 cells). **f**, PoSub neurons from freely-moving recordings grouped into three categories (FS, fast spiking; HD, head direction; slow non-HD), based on baseline firing rate, spike waveform (trough to peak) and head direction information (bits per spike). **g**, Cell type distributions for negatively and positively modulated cells. **h**, Example raster plots (*left*) and mean ± s.e.m. firing rate (*right*) of an example PoSub HD cell in awake conditions to Object, ScrambledD and ScrambledT stimuli. **i**, Same as (g), but for an example FS neuron. **j**, VOSIT and VOSID values for different types of positively modulated PoSub neurons (mean ± s.e.m.), p (VOSID, from left to right) = 0.032, 0.555, 0.017, p (VOSIT, from left to right) = 0.003, 0.985, 0.047, B-H corrected, Wilcoxon signed rank test (n (from left to right) = 58, 14, 57 cells).

First, we examined the nature of light-evoked activity in PoSub. We found that visual stimulation increased the firing rate of some PoSub neurons, whereas it decreased the firing rate of other neurons, a phenomenon we did not observe in anesthetized animals (**Fig. 3b,c**; also, some temporal properties of the light response also changed between awake and anesthetized conditions; **Extended data Fig. 9**). If we split PoSub neurons into positively and negatively visually modulated cells, we found that a preference for visual objects was specific to positively visually modulated cells (**Fig. 3d,e**).

Next, using the spike waveform, baseline firing rate, and head direction information encoded by each cell, we classified PoSub neurons into three cell types: HD cells, fast-spiking (FS) inhibitory cells and slow non-HD cells (**Fig. 3f**), as done previously^39^. We found that positively and negatively modulated cells were present in each group, however the majority of negatively modulated cells were HD cells (**Fig. 3g**). Furthermore, focusing solely on positively modulated cells, we found that a preference for visual objects (i.e. positive Object-Scrambled_T_ and positive Object-Scrambled_D_ responses) was present in HD cells and FS cells, but not slow non-HD cells (**Fig. 3h-j**).

### Visual objects dynamically refine population-level encoding of head direction

We next focused on PoSub HD cells to understand what was controlling the differential effects of visual stimulation on the firing rate of HD cells. We examined if there was population-level organization of visual responses. For a given animal, from freely moving recordings, we calculated the preferred firing direction of all simultaneously recorded HD cells and positioned each HD cell on a ring, with the angular position of each cell corresponding to its preferred firing direction (**Fig. 4a**). Next, for the same HD cells, from the head-fixed recordings we calculated each cell’s visual response (positive values indicate that visual stimulation increased the firing rate; negative values indicate that visual stimulation decreased the firing rate; **Fig. 4a**). We found that HD cells with similar preferred firing directions in the open field exhibited similar visual responses in the head-fixed experiment (**Fig. 4a**). In other words, HD cells that were positively modulated by visual stimulation had similar preferred firing directions, and HD cells that were negatively modulated by visual stimulation had similar preferred firing directions (**Fig. 4a**). To quantify this effect, for the ring generated for each mouse we computed the average preferred firing direction of positively visually responding HD cells and aligned this to 0° (see Methods; **Fig. 4b,c**). We found a correlation between visual response and angular position on the ring following alignment to 0° (**Fig. 4c**, R=0.294 (circular correlation), and **Extended Data Fig. 10**, R=0.316 (linear correlation)) that significantly differed from chance (**Extended Data Fig. 10**). Similarly, we found that HD cells whose preferred firing directions pointed towards 0° (preferred firing direction of 0° ± 45°) exhibited stronger visual responses than the remaining HD cells (preferred firing direction of 180° ± 135°; **Fig. 4d**). Thus, visual stimuli differentially modulate the firing rate of HD cells in PoSub as a function of preferred firing direction.

**Fig 4.**
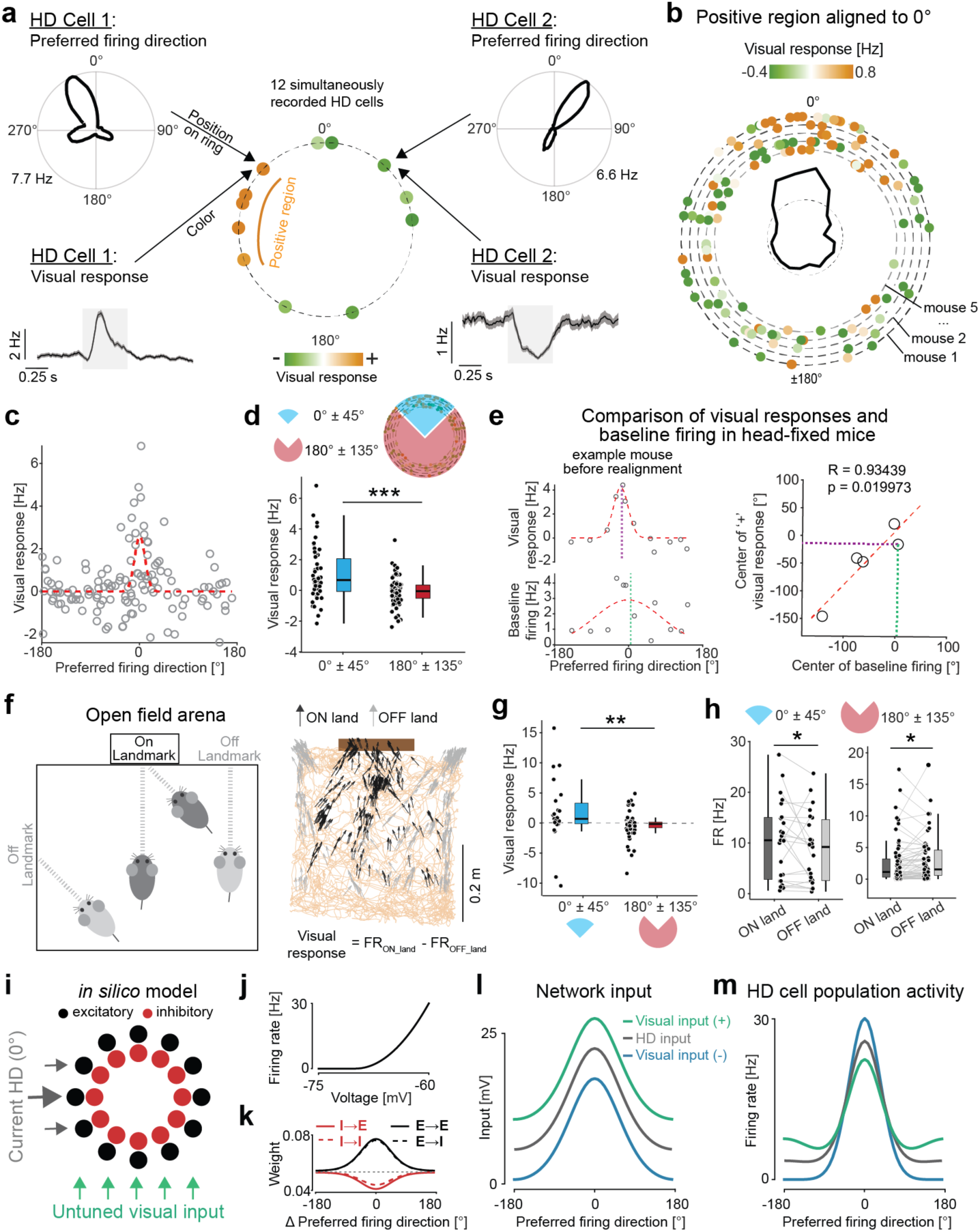
Visual objects dynamically refine population-level HD encoding. **a**, Schematic outlining how, for each mouse, we compared the visual response and preferred firing direction for all simultaneously recorded HD cells. In brief, the preferred firing direction is indicated by the angular position of each cell (dot) on the ring, and the visual response is indicated by the dot color (orange, positive; green, negative). Data here is from one example mouse. **b**, The rings (dashed circles with colored dots) are shown for 5 mice, with the center of the positive visual response regions aligned to 0°. Central ring: Dashed line represents mean visual response across all bins and bold line the deviations from it (20 angular bins). **c**, Scatterplot of HD cell visual response as a function of preferred firing direction following alignment to 0° (n = 127 cells from 5 animals). **d**, Comparison between the visual responses of HD cells with preferred firing directions 0° ± 45° (blue) or 180 ± 135° (red), p = 1.13*10^-4^, Mann-Whitney U-test (n = 51 and 76 cells). **e**, *left*, For an example mouse, the visual response as a function of HD preferred firing direction is compared to the baseline firing rate (in head-fixed) as a function of HD preferred firing direction (before realignment to 0°). Violet and green dotted lines represent the circular mean for the example mouse. **e**, *right*, For each mouse, the center of positive visual responses is plotted against the center of region with the highest baseline firing (n = 5 mice). **f**, *left*, Schematic illustrating how for HDs of 0° ± 45° (with the direction of the wall containing the visual landmark set as 0°), from some positions the mice look directly at the landmark (On landmark) whereas from other positions mice do not directly look at the landmark (Off landmark). **f**, *right*, For an example mouse, its location over time is shown (orange) with HDs within the range of 0° ± 45° colored by whether the mouse is On landmark (black arrows) or Off landmark (grey arrows). The visual cue is represented by the thick brown line. **g**, Visual responses in freely-moving conditions for HD cells with preferred firing directions within the range of 0° ± 45 ° and 180° ± 135°, p = 0.002, Mann-Whitney U-test (n = 23, 68 cells). **h**, For HD cells with preferred firing directions within the range of 0° ± 45 ° and 180° ± 135°, the firing rate for both On and Off landmark positions, p (from left to right) = 0.037, 0.016, Wilcoxon signed rank test (n (from left to right) =23, 68 cells). **i,** Supralinear stabilized ring network. Excitatory and inhibitory neurons are assigned a preferred head direction at which they receive the strongest tuned input (current HD input). In addition, all neurons receive an untuned input (visual input). **j-k,**Neurons in the network have a supralinear activation function (**j**), and their local weights are determined by the difference in their preferred firing direction (**k**). Weight and activation function parameters were set following Hennequin et al.^44^ and were varied to verify model robustness (see **Extended Data Fig. 11**). **l, m,** Increasing/decreasing the level of untuned input (**l**) has a differential effect on model HD cell firing rates (**m**), depending if the preferred head direction is in-field (current HD: 0 degrees), or out-field (180 degrees).

However, because the HD tuning curves above were calculated in one environment and the head-fixed recordings were performed in a different environment, it was unclear how the above-described visual responses related to the animal’s head-fixed HD. To answer this question, we took advantage of existing knowledge about population responses in the HD system. The HD system exhibits several hallmarks of a continuous ring attractor^40^: only HD cells with similar preferred firing directions—those encoding the mouse’s current head direction—can be active at a given time, resulting in a localized ‘bump of activity’ in the ring^5,41^. Furthermore, the angular offset between HD cells is maintained across environments^42^, making it possible to compare responses of similarly tuned neurons between different conditions. Thus, we tested whether the skewed distribution of positively and negatively visually responsive HD cells around 360° related to the ‘bump of activity’ in the HD ring attractor. Since HD cells continue to fire even when an animal is stationary^2,43^, the ‘bump of activity’ in baseline firing (in the absence of visual stimulation) should indicate which HD cells are encoding the animal’s current (head-fixed) head direction. We calculated the ‘bump of activity’ from baseline firing rates during head-fixation, indicating the current head direction, and found that it closely aligned with the region of the ring that exhibited positive visual responses (**Fig. 4e**). In other words, HD cells with preferred firing directions that pointed towards the visual stimuli (which were presented directly in front of the animal) were positively modulated by visual stimulation, whereas HD cells coding other directions tended to be unmodulated or negatively modulated by visual stimulation.

The visual stimulation experiments described above were performed in head-fixed conditions. Do visual objects similarly modulate HD cell firing in freely moving conditions? To test this, we set the direction of the wall of the open field arena that contained the visual landmark to be 0°. Next, we restricted our analysis to time points when the animal’s HD was within the range of 0° ± 45°, to focus on HDs over which it was possible for the mouse to directly face the visual landmark. Finally, we split the data into time points when the mouse was either directly facing the landmark (On landmark) or facing away from the landmark (Off landmark; **Fig. 4f**), while matching the distributions of head directions included in the two conditions (**Extended Data Fig. 10**). We then subtracted the average Off landmark firing rate from the average On landmark firing rate to obtain a visual response value in freely-moving conditions. We found that the visual response of HD cells with preferred firing directions aligned with the direction of the landmark (0° ± 45°) was significantly larger than the visual response of HD cells that coded for directions not aligned with the landmark (180° ± 135°; **Fig. 4g**). This arose because HD cells with preferred firing directions aligned with the direction of the landmark were positively modulated by the landmark, whereas responses of HD cells that coded for directions not aligned with the landmark were negatively modulated by the landmark (**Fig. 4h**). Together, these results show that visual landmarks dynamically modulate HD population activity as the animal navigates an environment, boosting the ‘bump of activity’ of the HD ring attractor coding the current HD when the animal is facing the landmark, while simultaneously decreasing activity on the ring away from the ‘bump of activity’.

Lastly, we asked whether our finding of a differential effect of visual input on HD cell firing as a function of preferred firing direction could arise from an untuned input (i.e. the same visual input to all PoSub cells, regardless of preferred firing direction). Indeed, it seems implausible that the visual input reaching PoSub would already be tuned by HD. We reasoned that a supralinear stabilized network^44^ (SSN) might recapitulate our observation seeing as 1) the SSN has previously accounted for tuning-specific experimental results when model units were organized in a ring architecture^44,45^ and 2) the SSN activity properties differ depending on the strength of the input^44^. This is due an architecture that relies on recurrent inhibition to stabilize the effects of strong recurrent excitation and a supralinear activation function, in which gain increases as stronger inputs are received (**Fig. 4i-k**). Here, the units of the SSN were provided with a spatially localized bump of activity, modeling activity within the HD ring attractor representing the current HD (**Fig. 4i-k**). Next, we examined the effect of the visual input to PoSub, modeled as an untuned input to the ring SSN network (**Fig. 4l**). We found that, despite the visual input to all units being the same, it evoked differential effects depending on each model unit’s angular distance from the current HD (**Fig. 4m**). Notably, the model recapitulated the experimental data, with visual input producing an increase in firing of model units coding for the animal’s current HD (represented by the location of the bump of activity in the ring), and a decrease in firing of model units coding HDs away from the current HD. Intriguingly, this effect arose only when visual stimulation corresponded to a decrease in global input to the SSN. These results were robust to parameter variations (**Extended Data Fig. 11**).

## Discussion

Combining brain-wide functional ultrasound imaging and both head-fixed and freely-moving electrophysiological recordings in mice, we made two central findings related to visual objects and the brain’s spatial navigation system. First, at the level of mean activity in brain areas, the spatial navigation system exhibits a preference for visual objects over scrambled versions of the same images. Second, visual objects directly modulate head direction coding in PoSub: a visual landmark boosts the firing rate of HD cells with preferred firing directions pointing toward the visual landmark, whereas it decreases the firing rate of HD cells that encode other HDs.

Interestingly, unlike in humans and non-human primates^26,27^, our brain-wide screen did not reveal that ventral visual cortical areas were more strongly activated by objects compared to scrambled images. While it is possible that spatial resolution limitations imposed by volumetric fUS (∼250 µm) could impede our ability to detect visual object preferences in particularly small brain areas, our findings are consistent with single cell calcium imaging experiments from mouse ventral stream areas that, to date, have failed to reveal individual neurons with preferred stimuli that resemble objects^20^ (though a slight preference for textures may exist in certain rodent ventral areas^19,46^). However, the lack of a clear preference for visual objects in the rodent ventral pathway does not mean that neurons in these areas are not involved in encoding visual objects. Indeed, previous studies in rodents trained on invariant visual object recognition tasks found that the amount of information contained about object identity increased along the ventral stream hierarchy^17,18^. Furthermore, in mouse V1, previous work has shown that while phase scrambling natural images does not affect mean population activity or response variance, it does affects higher- order correlations between neurons^42^. Thus, while mouse visual cortex appears capable of encoding information about visual objects and passing this information onto the spatial navigation system, unlike primates there may not be specific visual cortical areas dedicated to encoding specific classes of visual objects. Interestingly, beyond a potential role in visual object processing, it has been hypothesized that the rodent ventral visual stream may be important for visually-guided spatial navigation, as ventral visual cortical areas mostly encode regions of the visual field directly in front of the mouse, which could help them process visual landmarks^32^.

Our experiments demonstrate that several regions within the spatial navigation system, notably dorsal RSC and PoSub, preferentially respond to visual objects. Both of these areas are known to receive substantial input from the visual system and be involved with processing of landmarks^14,47,48^. RSC is critical for aligning reference frames, translating egocentric signals— such as viewpoint-specific visual inputs—into allocentric information, including the HD signal^48,49^. The RSC is thought to detect visual landmarks in the environment and integrate them with subcortical HD inputs, which are primarily driven by vestibular and proprioceptive signals, anchoring the HD signal to the external world^37,50–54^. PoSub is the cortical hub of the HD system^55^ and updates the HD signal with visual orientation cues^9,56^. It broadcasts the HD signal to downstream navigation structures such as the medial entorhinal cortex and hippocampus^57^. While the head-direction signal is ubiquitous in the medial entorhinal cortex^58,59^, hippocampal neurons also show tuning to head-direction in the presence of salient visual cues^60^. Furthermore, while our study mainly focused on the PoSub and RSC, we found other areas of the spatial navigation system that were modulated by objects, and it is possible that visual objects impact HD coding in those other areas as well. For example, our fUS screen identified the parasubiculum as preferring visual objects, and this area is known to play a role in conveying a directional signal to spatially tuned cells of the entorhinal cortex^61^.

As noted above, the preference for visual objects in the spatial navigation system that we describe does not appear to result from highly specific responses to individual objects by individual neurons. Instead, it appears to reflect a general preference for objects over scrambled images, which could arise from a preference for natural image statistics present in objects. This object preference that we describe could be useful for isolating objects from a noisy, visually cluttered environment and facilitate the use of visual objects as spatial landmarks. This preference for visual objects was present in both awake and anesthetized conditions, similar to what has been found for visual feature selectivity in IT cortex of primates^62^. Furthermore, seeing as the preference for visual objects was maintained in anesthetized conditions, it indicates that the response preference in spatial navigation areas represents a sensory, not behaviorally, driven phenomenon. Next, an interesting feature of our computational model of the HD system in PoSub is that it suggests that net input to PoSub decreases when the animal faces a visual cue, which in turn leads to increased firing of HD cells coding the current HD, but future work is required to test this model prediction. Lastly, while the anatomical connectivity between primary visual cortex, higher visual areas, and RSC and PoSub is well mapped out^48,47,63–65^, the specific pathways and computations that enable visual objects to get encoded and modulate the spatial navigation system remain to be elucidated.

Our findings in PoSub show that the firing rate of HD cells coding the current head direction increases when an animal is facing a visual object. At the same time, the firing rate decreases for HD cells that are not coding the current head direction. This indicates that in moments when the visual landmark is in the centre of the mouse’s visual field, the certainty about the current heading direction is increased. However, whether this visual object-mediated increase in gain is specific to the HD system, or also extends to other types of spatially tuned neurons throughout the spatial navigation system remains an open question.

## Methods

### Animals

Experiments performed in Germany complied with the institutional guidelines of the Max Planck Society and were approved by the local government (Regierung von Oberbayern). Experiments performed in Canada were done in accordance with the Canadian Council on Animal Care and approved by the Montreal Neurological Institute’s Animal Care Committee. Male and female C57BL/6 mice aged 8-12 weeks were used in all experiments. They were housed in groups under a 12-hour light-dark cycle with unrestricted access to standard diet and water.

### Cranial Window Surgery

Surgeries were performed as described previously^30^. Briefly, mice were anesthetized with a subcutaneous injection of fentanyl (0.05 mg/kg), midazolam (5 mg/kg), and medetomidine (0.5 mg/kg) (FMM cocktail). They were secured using a bite bar and kept at 37°C with a temperature controller (Supertech). To prevent their eyes from drying, a hydration gel (Bayer, Bepanthen) was applied. A large cranial window was created in the skull using a dental drill, extending from bregma +2.25 mm AP to bregma -4.00 mm AP, while ensuring the dura remained intact. A pre-prepared COMBO window^30^ was adhered to the exposed bone using cyanoacrylate glue (Pattex) and the contact was reinforced with dental cement (Super-Bond). Anesthesia was reversed after surgery by administering subcutaneous flumazenil (0.5 mg/kg) and atipamezole (2.5 mg/kg). Postoperative analgesia was managed with subcutaneous injections of buprenorphine (0.1 mg/kg). After a 7-day recovery period, the FMM cocktail was administered again, and a head plate was attached to the COMBO window. Mice were allowed at least 3 days of additional recovery before being gradually acclimated to the experimenter, the behavioral setups, and tasks.

### Habituation to Head Fixation

For awake fUS recording sessions, all animals underwent gradual habituation to a head-fixed context. For the first 1 to 2 days, they were allowed to explore the behavior rig. Over the next 5+ days, the duration of head fixation increased daily from 10 minutes to 60 minutes.

### fUS Acquisition

fUS-imaging data was collected using a 32 × 32 channel matrix probe (15 MHz, 1024 total elements, spatial resolution: 220 × 280 × 175 μm³, Vermon) connected to a Vantage 256 system (Verasonics, Inc.) and controlled by a custom volumetric fUS acquisition module (AUTC). A 4× multiplexer linked the 1024 channel probe to the 256-channel system, with the beamforming and sequences adjusted accordingly. For fUS recordings under anesthesia, mice were injected with the FMM cocktail and kept at 37° with a temperature controller. Next, the mouse was head-fixed in a holding tube and the matrix probe was positioned to cover the entire cranial window using a three-way translation stage. A compound ultrasound image was created from the summation of plane wave emissions at -4.5, -3, -1.5, 0, 1.5, 3, and 4.5 degrees. A power Doppler image was generated from the incoherent average of 160 compound ultrasound images acquired at 400 Hz. Clutter filtering was performed in real time by decomposing the ultrasound stack using singular value decomposition, removing the first 20% of singular vectors. This resulted in a power Doppler image approximately every 500 ms.

### Electrophysiology Recordings under anesthesia

Data was recorded at 20 kHz using RHX software from 32-channel silicon probes (Cambridge NeuroTech, type H10b) connected via a SPI interface cable (Intan technologies, C3216) to a USB interface board (Intan technologies, C3100). Mice were anesthetized with the FMM cocktail and head-fixed in a holding tube, aligned to the probe manipulator’s coordinate system. Small perforations (0.5 mm diameter) were made above the region of interest, and the probe was lowered into the brain at 2 μm/s. Recordings began 10 minutes after reaching the target depth. For separate recordings on different days, perforations were sealed with Kwik-Cast silicone sealant. CM-DiI (Thermo Fisher, CellTracker) was applied to the probe before brain insertion to identify recording sites post hoc.

### Silicon probe implantation for chronic (awake) electrophysiology

Animals were anesthetized with 5% isoflurane and positioned in a stereotaxic apparatus. Anesthesia during the surgery was maintained with 1.5-2% isoflurane. Animals were given a subcutaneous saline injection (∼0.8 ml) and carprofen (20mg/kg), as well as a subcutaneous 1:1 lidocaine-bupivacaine mixture underneath the incision site as soon as they were placed in the stereotaxic apparatus.

A thin layer of light-cured adhesive (Kerr OptiBond) was applied to the skull to help implants better adhere to the skull. A head-fixation bar was cemented to the skull using a dental acrylic cement (Unifast Trad). A silver wire was then implanted into the cerebellum to serve as a reference electrode. Two types of silicon probes were utilized for the experiments: a single-shank probe (Cambridge NeuroTech H5) or a single shank probe with a Molex connector (Cambridge NeuroTech H5). Both probes were mounted on a moveable microdrive.

A craniotomy was performed above the target area, creating a hole slightly larger than the probe itself, approximately 0.1 mm wider on each side. The silicon probes were implanted to the following coordinates: AP (from Lambda) +0.30, ML -2.35, DV -1.10. The probe was carefully lowered into position and affixed to the skull using dental acrylic cement (Unifast Trad). To minimize electrical interference during recordings, a copper mesh was attached to the skull surrounding the implant using a light-cured flowable composite (Fusion Flo).

After surgery, animals were given a second subcutaneous saline injection (∼0.8 ml) and placed on a heated pad until fully recovered. Post-surgery, animals were housed individually and received daily carprofen injections for at least three days. Animals were allowed to recover for 7 days post-surgery before further experimentation.

### Electrophysiological recordings in awake animals

The probe was lowered slowly into the postsubiculum using the microdrive over the course of 2- 3 hours. Brain tissue was allowed to recover for 2 hours before the start of the recording. Neurophysiological signals were recorded continuously at 20 kHz using a 256-channel RHD USB Interface board (Intan Technologies) and captured with Intan RHX software.

Behaviour of the mouse during freely-moving open field arena sessions was tracked using reflective markers that were mounted on the copper mesh or pre-amplifier. These were tracked in 3D with infrared cameras and Motive 2.0 motion capture system (Optitrack). Video was captured by an additional camera mounted overhead. Behavioural tracking was acquired at a rate of 120 Hz and synchronized with the electrophysiological signal using voltage pulses registered by the RHD USB Interface Board. The mice were allowed to roam freely in a circular or square environment for 15-20 minutes. Following the freely-moving session, animals were immediately head-fixed and presented with the visual stimulation task.

### Visual stimulation

Images were presented at a size of 60° with the center of the image at 0° azimuth and +20° elevation on a 61 cm computer monitor (Dell, U2415b) using the PsychoPy toolbox. The monitor was placed 18 cm in front of the mouse. For fUS experiments, we employed a block design consisting of 12 s of gray followed by 12 s of images (12 images, each presented for 0.5s interleaved with 0.5s gray, with images presented in random order across trials). Each recording run consisted of eight repetitions of eight blocks (four object blocks and four texture scrambling blocks) in random order. For object presentation in electrophysiological experiments, images (48 objects, 48 textures, 48 diffeomorphic transformations) were presented in pseudorandom order for ten times for 0.5s and interleaved with at least 0.5s of gray.

### Image preparation

Original images were resized to 1024x1024 pixels and put on a medium gray background. For each of these images we created a control image with matching statistics using the described texture synthesis algorithm^33^ with 25 iterations. Subsequently, all images were gamma corrected and the SHINE toolbox^66^ was used to luminance match all images. Finally, to avoid edge effects a small border (7 pixels around the image) was blurred using a gaussian filter. For stimuli used in electrophysiological experiments we additionally created a second set of control images (diffeomorphic transformations) using a previously published algorithm^38^. Additionally, to avoid potential changes in spatial frequency or luminance content introduced by this procedure, for electrophysiology experiments with two scrambling methods, the whole amplitude spectra (specMatch.m) as well as the luminance histogram (histMatch.m) of all images and their respective controls were equated in ten iterations using the SHINE toolbox.

### Analysis fUS

#### fUS Preprocessing

The fUS time series was first registered to the Allen Brain Institute’s mouse brain atlas. For this, first all sessions of a mouse were aligned using an automated approach based on the imregtform.m function. Then, a high-resolution power Doppler image was created by averaging 100 fUS frames from a single session, which was then manually registered to the atlas using anatomical landmarks to create a transformation matrix using a previously published toolbox^25^. This matrix was applied to data from other sessions of the same mouse. The fUS time series was preprocessed using custom MATLAB scripts on a voxel-by-voxel basis. Temporal interpolation was performed to achieve a constant frame rate of 2 Hz. The relative change in power Doppler signal was calculated by subtracting the baseline signal from each time point and dividing by the baseline signal. Slow drifts were removed using a fifth-order high-pass Butterworth filter with a cutoff frequency of 0.056 Hz. To eliminate movement artifacts during awake experiments, the top 10% of temporal principal components extracted from non-brain voxels was regressed out from all voxels.

### fUS activation

To assess brain activation at the voxel-wise level, the preprocessed fUS data was temporally smoothed (four frames) and fitted using a General Linear Model (GLM) using the MATLAB function glmfit.m. Model regressors included the visual stimulation block stimuli, convolved with a single-gamma hemodynamic response function. The resulting T-scores where averaged across animals/sessions and plotted on top of the Allen Brain Atlas (thresholded by p-values, see below).

To assess activation at the brain region level, first the preprocessed and trial-averaged (only trials of the same type of stimulus; objects or texture) whole-brain data was segmented into individual brain regions. For this, anatomical regions from the Allen reference brain atlas were consolidated into 100 brain regions and the data from all voxels within each region was averaged. Next, the correlation coefficient between the stimulus timing and the fUS signal of each brain region was calculated. To not remove regions that only respond to object or texture blocks respectively, the correlation was performed three times (objects only, scrambled only and both combined). Regions with significant p-values (see below) for one or more of the three correlations were defined as active.

Similarly, to find brain regions with differences between object and scrambled stimuli, a preference index for each region (or voxel) was calculated on the preprocessed and trial-averaged fUS data. For this, the area under the curve during the response window for texture stimuli was subtracted from the response to object stimuli and normalized by the sum of the two responses (VOSI_T_). To unbiasedly define a response window for each region/voxel, we averaged all data of a given experiment (sessions, animals, stimuli, all trials) to end up with a single response kernel. The start of the response window was defined as the point with maximum rate of change. The end of the response was defined as the point where the response dropped below 70% of the peak response (30% for awake fUS data due to different temporal dynamics). In case this method did not result in a meaningful response window (window very short or start/stop point before/after stimulus onset/offset), a generic stimulus window (from stimulus onset to stimulus offset) was used.

To determine significance, a mixed effects model was fitted on T-scores, correlation scores or preference indices for each voxel/region using the fitlme.m function. Resulting p values were corrected for multiple comparisons using False Discovery Rate (FDR)-correction.

### Region and voxel clustering

Preprocessed, segmented and trial-averaged (trials of the same stimulus type) fUS data of visually active regions were clustered using a MATLAB UMAP^67^ implementation (Meehan, Version 4.4; with correlation as distance metric and the following parameters: n_neighbours=5, cluster_detail=’very low’). Similarly, for hierarchical clustering, the segmented and trial-averaged fUS data of visually active regions was clustered using the MATLAB function dendrogram.m (distance metric = correlation, linkage method = weighted). For clustering at the level of voxels, preprocessed and trial-averaged fUS data of visually active voxels (defined by the above described GLM approach) were clustered using k-means with correlation as distance metric and number of clusters defined by the silhouette method.

### Analysis electrophysiology Preprocessing

Electrophysiological recordings were processed with Kilosort3, and the spike sorting results were manually refined using the Phy software (https://github.com/cortex-lab/phy). Only units that exhibited stable responses throughout the session and achieved high Kilosort quality scores were included in the analysis. To estimate continuous firing rates, single-unit responses were binned with a 10 ms bin size. Except for raster plots, the ten trials of the same image were averaged. Raster plots were plotted using the Gramm toolbox^68^.

### Visual object and scrambled stimuli

To select only visually responsive units, we averaged all trials and stimuli of a cell, temporally smoothed the response (10ms window) and determined the response amplitude as the difference of the peak of this overall averaged response minus the baseline firing rate. Cells with an amplitude below 1Hz were excluded from the analysis (in awake electrophysiology all cells were included). To determine the difference between object and control stimulus presentation, the average firing rate during the response window (stimulus onset to stimulus offset) was calculated and the median response to control stimuli (Scrambled_T_ or Scrambled_D_) was subtracted from the median response to object.

### Receptive field mapping

White squares (30° x 30° visual angle) were flashed at 15 different positions (3 elevation positions x 5 azimuth positions) of the screen at random order. Each position was repeated 20 times. The stimulus was presented for 0.5s followed by at least 0.5s of gray. Only units that had a response amplitude larger than 1 Hz for at least one of the screen positions were included in the analysis. To determine how many positions of the screen resulted in a significant activation, we tested at each position whether the firing rates during the response (40ms around peak response) was significantly different from the firing rates before and after stimulus presentation across the 20 repetitions of the stimulus (Wilcoxon signed rank test, B-H corrected).

### Spatial frequency tuning

Black and white striped stimuli of different spatial frequencies and two orientations (horizontal and vertical) were presented at random order. Like object stimuli, the area of these stimuli spanned 60° visual angle and were centered at 0° azimuth and +20° elevation, and were displayed for 0.5s, interleaved with 0.5 s full-field gray. Each stimulus was shown 20 times. Only units with a significant response to at least one stimulus (Wilcoxon signed rank test, FDR corrected) were included in the analysis. For each cell and spatial frequency, the maximum firing rate during the response window in response to the more preferred orientation was calculated. This resulted in a per cell spatial frequency tuning curve (the maximum represents the preferred spatial frequency) that, for visualization of units with different firing rates, was rescaled between zero and one.

### Chirp stimulus

A full-field temporal chirp stimulus was presented 20 times, interleaved with ten seconds of full-field gray. The stimulus intensity smoothly transited from black to white with increasing frequency from around 0.3Hz up to 4Hz. Only units with a response amplitude larger than 3 Hz were included for analysis. To determine when a cell stopped responding to the chirp stimulus, first, the normalized firing rates of each cell were decomposed in time-frequency using 1-D wavelet transformation (cwt.m function). Then, the power across all frequencies was summed and the point in time during the stimulus where the power dropped below 1.5 times of the pre-stimulus power was calculated. The last chirp frequency presented before this time point represents the maximal frequency the cell responded to.

### HD tuning metrics

HD tuning curves were calculated as previously described^39^. Briefly, markers on the animal’s headcap were tracked and connected to a three-dimensional polygon allowing calculation of head direction as the horizontal orientation (yaw) of the polygon. The yaw was measured in global coordinates (i.e., aligned with the environment’s axes, which remained consistent throughout the study). HD tuning curves were generated by taking the ratio of histograms of spike counts to the total time spent in each direction, using 1° bins and smoothing with a Gaussian kernel (3° standard deviation). The preferred firing direction of a cell is defined as the mean direction of the tuning curve.

To determine how strongly a cell is tuned to head direction, for each cell we determined the head direction information as previously described^39^ with this formula:

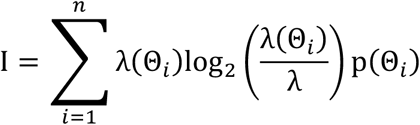

Here, n represents the total number of angular bins and λ(Θ_i_) represents the firing rate of the cell in the ith angular bin, λ denotes the neuron’s overall average firing rate during exploration, and p(Θ_i_) indicates the occupancy (normalized time spent) in direction Θ_i_. The information rate (I, expressed in bits per second) was then adjusted by the neuron’s average firing rate to yield an information content metric (head direction information, measured in bits per spike).

### Cell type classification

PoSub cells were classified as previously described^39^. Briefly, cells with a baseline firing rate larger than 10 Hz (during freely moving condition) and a trough to peak duration shorter than 0.4 ms were classified as fast-spiking neurons. In contrast, cells with baseline firing rates below 10 Hz and trough to peak duration above 0.4 Hz were considered slow-spiking neurons. This group was further subdivided into HD cells (head direction information above 0.1) or slow non-HD cells (head direction information below 0.1). All other cells were not classified.

### Visual responses in head-fixed conditions, and realigning HD based on visual responses

To determine the overall visual response of a cell, first all trials of the same images were averaged and then the median firing rate during the baseline period was subtracted from the median response to all images during the stimulus period. Cells with a visual response value above 0.2 Hz were assigned to the positively modulated group. Cells with a visual response smaller than -0.2 Hz were assigned to the negatively modulated group. To visualize the visual responses with respect to the cell’s HD direction tuning (activity bump, center ring in Fig. 4b), the realigned cells of all mice were binned into 20 angular bins based on their preferred head direction. Then the smoothed and binned visual responses were plotted as scaled deviation (0.1 scaling factor) from the average response (dashed line circle).

In each mouse we observed an accumulation of cells with similar head direction tuning and increased visual activity (and baseline activity), however the center of this activity was differently oriented for each mouse likely because there was no systematic remapping when moving the mice from the freely moving condition to the head-fixation setup. As such, we realigned the HD cells’ preferred directions such that the cluster of positively visually modulated HD cells was centered around 0°. For this, in each mouse the circular mean of the preferred directions of the cells that positively responded to the visual stimulus was determined, and this mean direction was subtracted from the preferred directions of all cells, concurrently shifting the whole ring. For shuffled distributions, the realignment process per mouse was repeated 100,000 times, with the visual responses of all cells of a mouse randomly shuffled and the preferred directions kept constant. To calculate the center of baseline firing in head-fixed condition, the circular mean of the cells with high baseline firing rate (> median of all cells of a mouse) was determined.

### Visual responses in freely moving mice

For this analysis, only recording sessions performed in a rectangular arena were considered (4 out 5 mice), so that the geometry was the same for all analyses (one session in a circular arena was excluded here). First, positional and head direction data was resampled to match 25 ms binning of the spiking data (resample.m). Then, to identify time points when the mouse was looking directly at the visual landmark (On landmark) or looking away from the landmark (Off landmark), at each spatial location we checked if and where the extension of the head direction immediately in front of the animal intersected with the north wall positions where the landmark was placed. Spatial locations and head directions with intersections on the landmark (but not on the edges: 1 cm buffer zone) were considered “On landmark”. Similarly, spatial locations and head directions with intersections beside the landmark were considered “Off landmark”. To ensure the intersections were “Off landmark”, intersections closer than 10cm from a landmark edge were excluded. To exclude positions and head directions with extreme cue viewing angles, which would result in a strong distortion of visual cue, in both cases, only time points, where the absolute head direction was 0° ± 45° were included. Moreover, to ensure that the selection of the time points did not result in a bias in selection of head directions (which could result in differences between “Off landmark” and “On landmark” due to HD tuning of the cells), we subsampled the larger of the two indices and matched the histogram of head directions for each session. Finally, we averaged the firing rates during “On landmark” and “Off landmark” time points and subtracted the two to get the visual response per cell.

### Response delay

For each HD cell, all trials and stimuli presentations were averaged to get an overall response of the cell. The response delay was defined as the point when the firing rate of the cell rose above two times the standard deviation of the pre-stimulus period.

### Perfusion and Histology

Following the termination of the electrophysiological experiments, animals were deeply anesthetized using sodium pentobarbital or ketamine (120 mg/kg) and xylazine (16 mg/kg) cocktail and perfused transcardially first with 0.9% phosphate-buffered saline solution followed by 4% paraformaldehyde solution.

In case of chronic recordings, the microdrive was retracted to remove the probe from the brain. Then brains were isolated and kept in a 4% paraformaldehyde solution for 24 hours, after which the solution was changed to 30% sucrose in phosphate-buffered saline until sinking. After freezing in a -80° freezer, brains were sectioned with a freezing microtome coronally in 40 μm slices. Sections were washed, counterstained with DAPI and mounted on glass slides with ProlongGold fluorescence antifade medium. Sections containing probe tracts were additionally incubated with a Cy3 anti-mouse secondary antibody (1:200 dilution; Cedarlane, 715-165-150) to help visualize the electrode tract. Widefield fluorescence microscope (Leica) was used to obtain images of sections and verify the tracks of silicon probe shanks.

In case of acute recordings, after perfusion, brains were collected and incubated in 4% PFA at 4°C overnight. Then brains were washed and embedded in 4% agarose. Coronal slices were cut using a vibratome (Leica microsystems, VT1000S) with a thickness of 100 μm and stained with DAPI (Thermo Fisher Scientific, D1306 or R37606). Brain slices were washed and mounted on microscope slides in Vectashield (Vector Laboratories, H-1200). Images were captured using a Leica TCS SP8 laser scanning confocal microscope with a 20x air objectives (Leica Microsystems). Images were processed using LAS X (Leica Microsystems) and Adobe Photoshop software.

### SSN model

The supralinear stabilized ring network model followed Hennequin et al.^45^ Specifically, the network contained 𝑁*_E_* = 50 excitatory and 𝑁*_I_*_’_ = 50 inhibitory neurons evenly spaced around a ring with angle 𝜃 ∈ [−𝜋, 𝜋]. The circuit dynamics followed

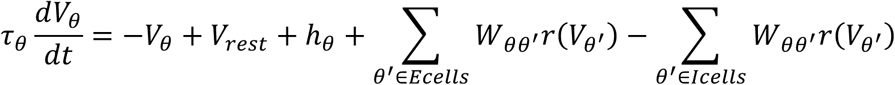

where 𝑉_𝜃_ and 𝜏_𝜃_ correspond to the membrane potential and time constant of a model unit at angle 𝜃, 𝑉*_rest_*, is the neuronal resting potential, ℎ_𝜃_ is the external input, and 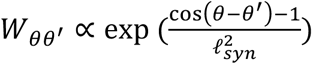 the strength of synaptic connection from neuron 𝜃^7^ to neuron 𝜃, which is further scaled such that the sum of incoming weights corresponds to 𝑊_EE_, 𝑊_IE_, −𝑊_EI_ and −𝑊_II_. The input was taken to be

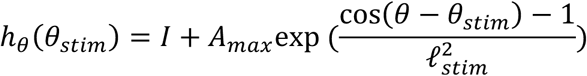

which consists of a tuned component at 𝜃*_stim_* = 0 with half width ℓ*_stim_* and magnitude 𝐴*_max_*, and an untuned component of magnitude 𝐼. We assumed that E and I model units at a given HD are driven equally strongly by the stimulus.

Finally, the firing rate of each cell is a threshold-power law function of its membrane potential:

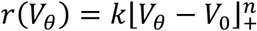

Parameters were taken as the default in Hennequin et al.^45^ (see **Supplementary Table 5**). To test the parameter robustness of our results, a population of 1250 networks with different weight hyperparameters was sampled from a uniform distribution (see **Supplementary Table 5** for bounds). For each of these networks, the change in rate of neurons aligned with the input HD (0°, corresponding to the current HD; ‘in-field’) and opposite to that HD (180°; ‘out-field’) was calculated for a small change in input (𝜕𝐼 = 0.1).

## Acknowledgements

We thank L. Mainville for performing histology for post hoc identification of electrode tracks. We thank T. Frank, A. Krishnaswamy, J-M. Martinez de Paz, and P. Wanken, T. Gollisch and N. Gogolla for helpful discussions and feedback on the manuscript. We thank G. Montaldo and A. Urban for technical support with fUS data acquisition. We thank H. Lin and A. Aliyeva for their contributions on aspects of this project not included in the manuscript. D.S. was supported by the Swiss National Science Foundation (SNSF) Early Postdoc.Mobility no. 194957, SNSF Postdoc.Mobility no. 211087 and Deutsche Forschungsgemeinschaft Walter-Benjamin Programm (Stelle) no. SI 2831/1-1. E. M. was funded by the Max Planck Society and the Deutsche Forschungsgemeinschaft (DFG, German Research Foundation) under Germany’s Excellence Strategy - EXC 2067/1- 390729940. We acknowledge funding from the Vanier Canadian Graduate Scholarship to S.S.C. Canada Research Chairs to A.P. and S.T.; Canadian Institutes of Health Research Project Grants (190289 and 180330) and National Sciences and Engineering Research Council Discovery Grant (RGPIN-2018-04600) to A.P.; Alfred P. Sloan Foundation Research Fellowship, Human Frontiers Science Program Career Development Award, Office of Naval Research Global Grant, and a Canadian Institutes of Health Research Project Grant (185933) to S.T.

## Author Contributions

Experiments were designed by D.S., S.T. and E.M. fUS experiments were performed by D.S. Anesthetized electrophysiological experiments were performed by D.S. and J.L.M. Awake electrophysiological experiments were performed by H.D. and S.S.C. Data was analyzed by D.S. SNN modelling work was performed by D.L. The manuscript was written by D.S., A.P., S.T. and E.M.

## Data and code availability

All data and code will be made available on public repositories upon publication.

## Extended Data Figures

**Extended Data Fig. 1.**
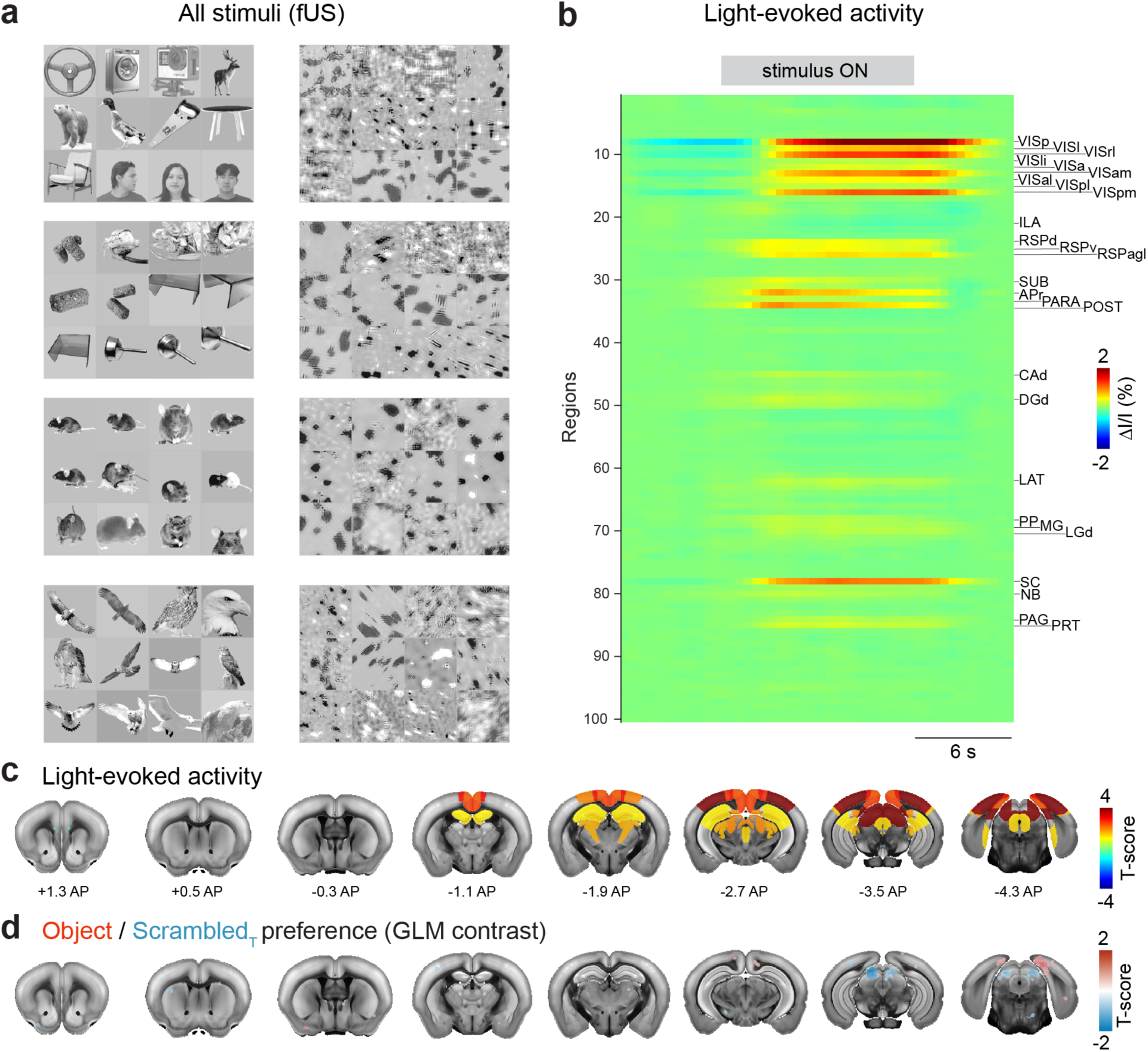
Brain-wide fUS imaging in anesthetized animals using object and control stimuli. **a**, All images used for visual stimulation in fUS experiments. Each block of images contains 12 object images (left side) or the same images scrambled using texture scrambling (right side). Blocks of images as well as images within blocks were presented pseudo-randomly. **b**, Area-segmented data shown as an average across all image blocks and sessions. Significant areas (correlation analysis, see methods) are indicated by area names (Allen Brain Atlas names), p < 0.05, FDR-corrected, mixed effects model (n = 56 sessions from 7 anesthetized animals). **c**, All significantly visually-responsive brain areas (as in b) color coded by the region-average T-score shown on coronal brain slices at indicated positions. **d**, GLM contrast between all Object and all ScrambledT blocks, p<0.01, uncorrected, mixed effects model (n = 56 sessions from 7 anesthetized animals).

**Extended Data Fig. 2.**
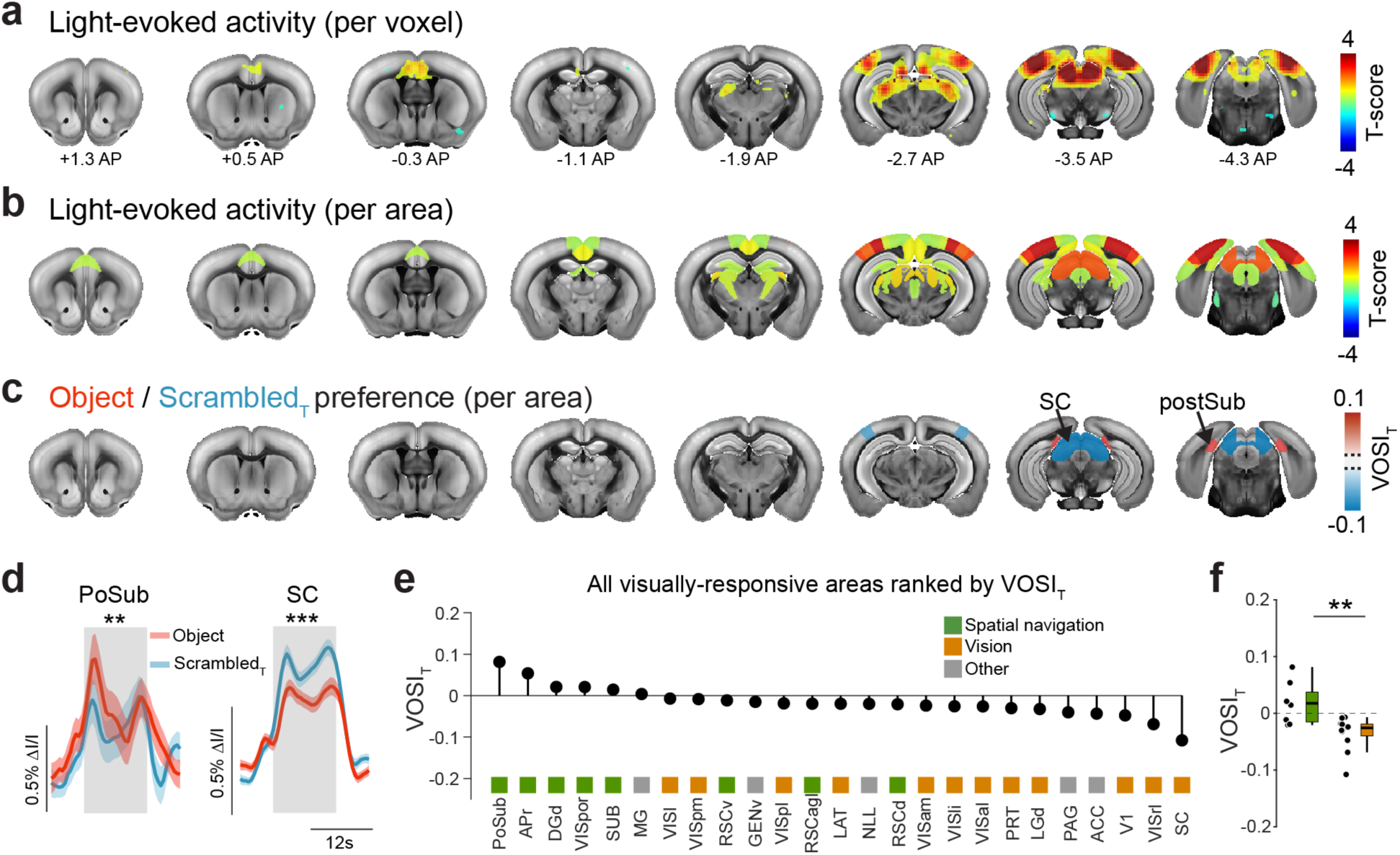
fUS imaging in awake animals is consistent with findings in anesthetized animals. **a**, Visually-evoked fUS responses (GLM, all image blocks as regressors) overlaid on coronal brain images at indicated positions. Only T-scores significantly different from zero are shown, p<0.05, FDR-corrected, mixed effects model (n = 83 sessions from 7 awake animals) **b**, All significantly visually-responsive brain areas (correlation analysis, see methods) color coded by the region-average T-score, p<0.05, FDR-corrected, mixed effects model (n = 83 sessions from 7 awake animals). **c**, Areas with a VOSIT significantly different from zero are colored by their VOSIT values. p<0.05, FDR-corrected, mixed effects model (n = 83 sessions from 7 awake animals) **d**, Session-averaged fUS responses (shaded area represents s.e.m.) to Objects and ScrambledT images from postsubiculum and SC. p(PoSub)=0.002, p(SC)=8.4*10^-4^, Bonferroni-Holm (B-H) corrected, mixed effects model (n = 83 sessions from 7 awake animals) **e**, Rank ordering of visually responsive brain areas according to the VOSIT values (*black dots*), color-coded (*squares*) according to whether they belong to Spatial navigation (*green*), Vision (*orange*) or Other (*grey*) brain networks. **f**, Comparison of VOSIT values for 8 spatial navigation areas (*green*) and 12 visual areas (*orange*) brain, p=0.003, Mann-Whitney U-test.

**Extended Data Fig. 3.**
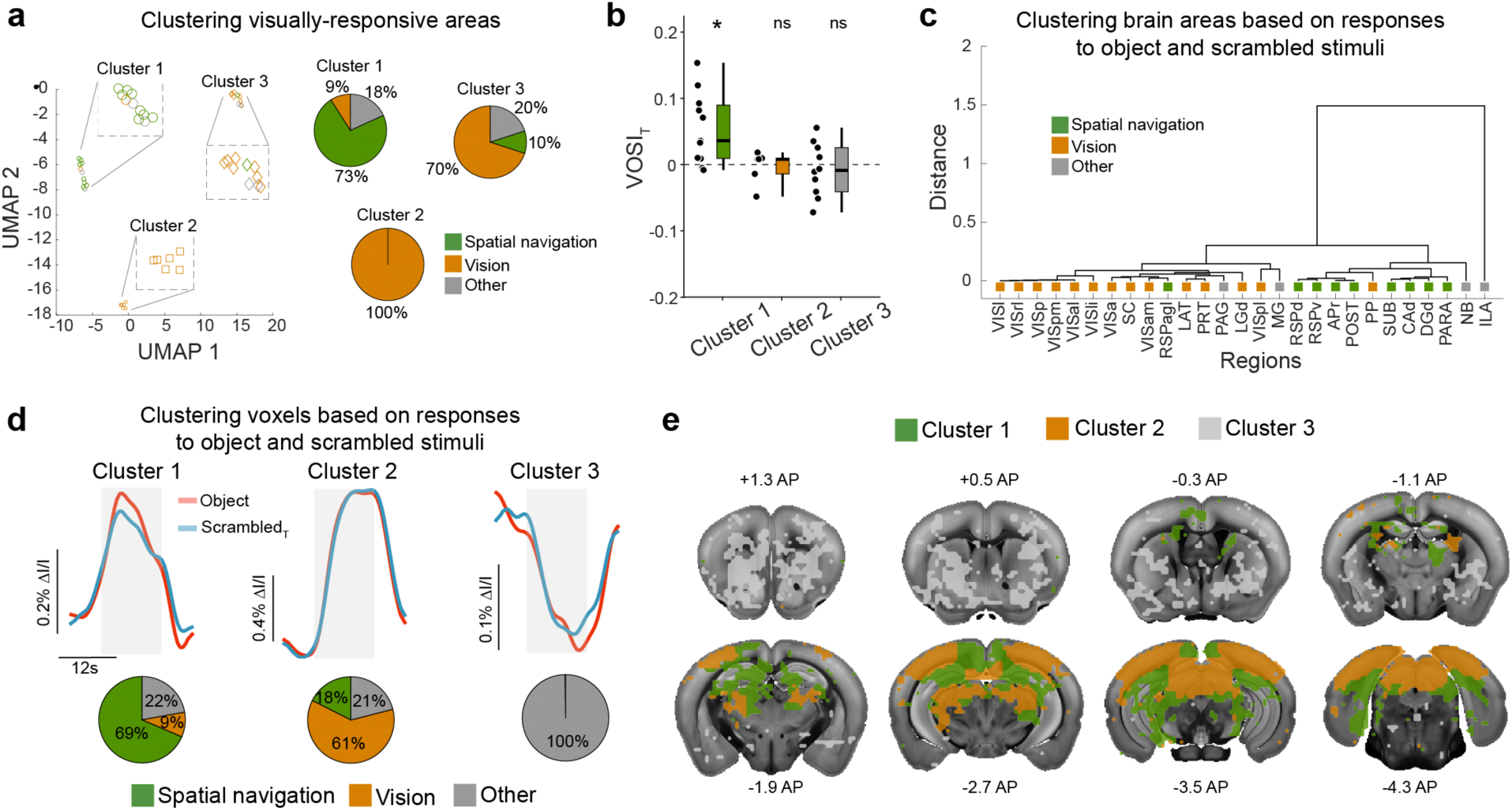
Unbiased clustering of brain regions, and voxels, groups spatial navigation areas as visual object preferring. **a**, UMAP-based clustering of the 27 visually active areas results in three clusters. Cluster 1 is dominated by regions of the spatial navigation system, while Clusters 2 and 3 mostly contain regions belonging to the visual system. **b**, Consistent with Figure 1, and the presence of many spatial navigation areas in Cluster 1, only regions of Cluster 1 display significant objects sensitivity, p (from left to right) = 0.014, 1.391, 1.391, B-H corrected, one-sample Wilcoxon signed rank test (n (from left to right) = 11, 6, 10). **c**, Similarly, hierarchical clustering of all visually-responsive areas separates spatial navigation areas from visual areas. **d**, K-means clustering of all visually-responsive voxels into three clusters results in one cluster (Cluster 1) mostly containing voxels belonging to Spatial navigation areas, another cluster (Cluster 2) dominated by voxels of Visual areas and the last cluster exclusively populated by voxels of Other brain areas. Time-resolved fUS traces show the average response for all voxels of indicated clusters separated by image type. **e**, Clusters as in (d) shown in the brain at indicated coronal positions.

**Extended Data Fig. 4.**
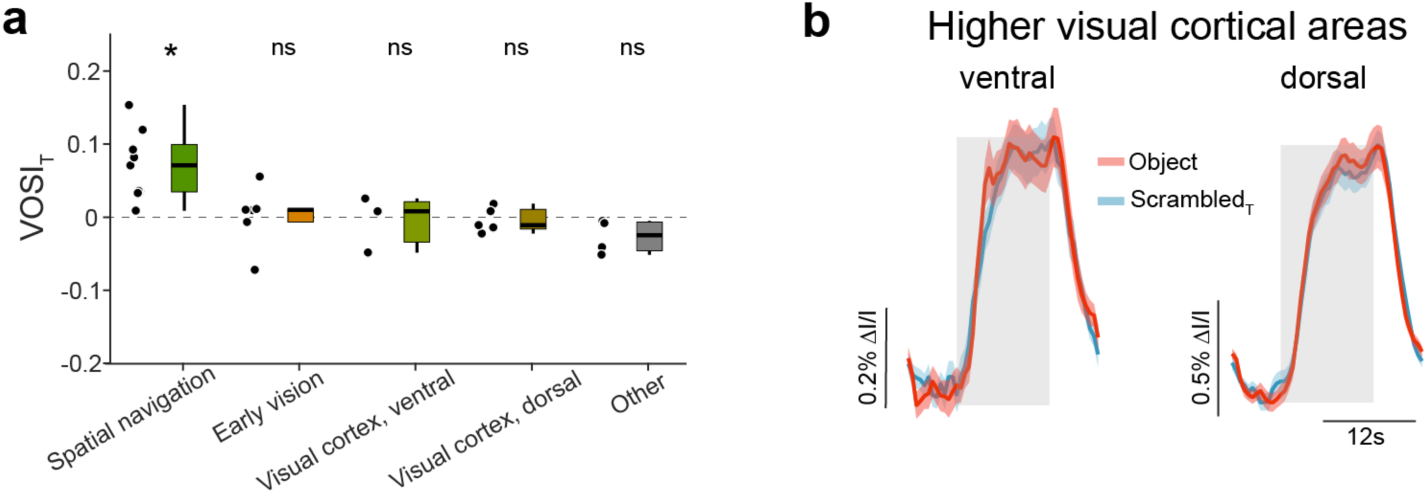
No evidence for a preference for object over scrambled images in dorsal or ventral visual cortical areas. **a**, VOSIT values as in Fig. 1h, with more refined grouping for “Vision” areas, p (from left to right) = 0.02, 1.688. 1.25, 1.688, 0.5, B-H corrected, one-sample Wilcoxon signed rank test (n (from left to right) = 9, 6, 3, 5, 4). **b**, Session-averaged fUS responses (mean ± s.e.m.) to Objects and ScrambledT averaged across all areas of indicated groups (ventral: VISl, VISli, VISpl; dorsal: VISrl, VISa, VISal, VISam, VISpm), p (form left to right) = 1.212, 1.212, B-H corrected, mixed effects model (n = 56 sessions from 7 anesthetized animals).

**Extended Data Fig. 5.**
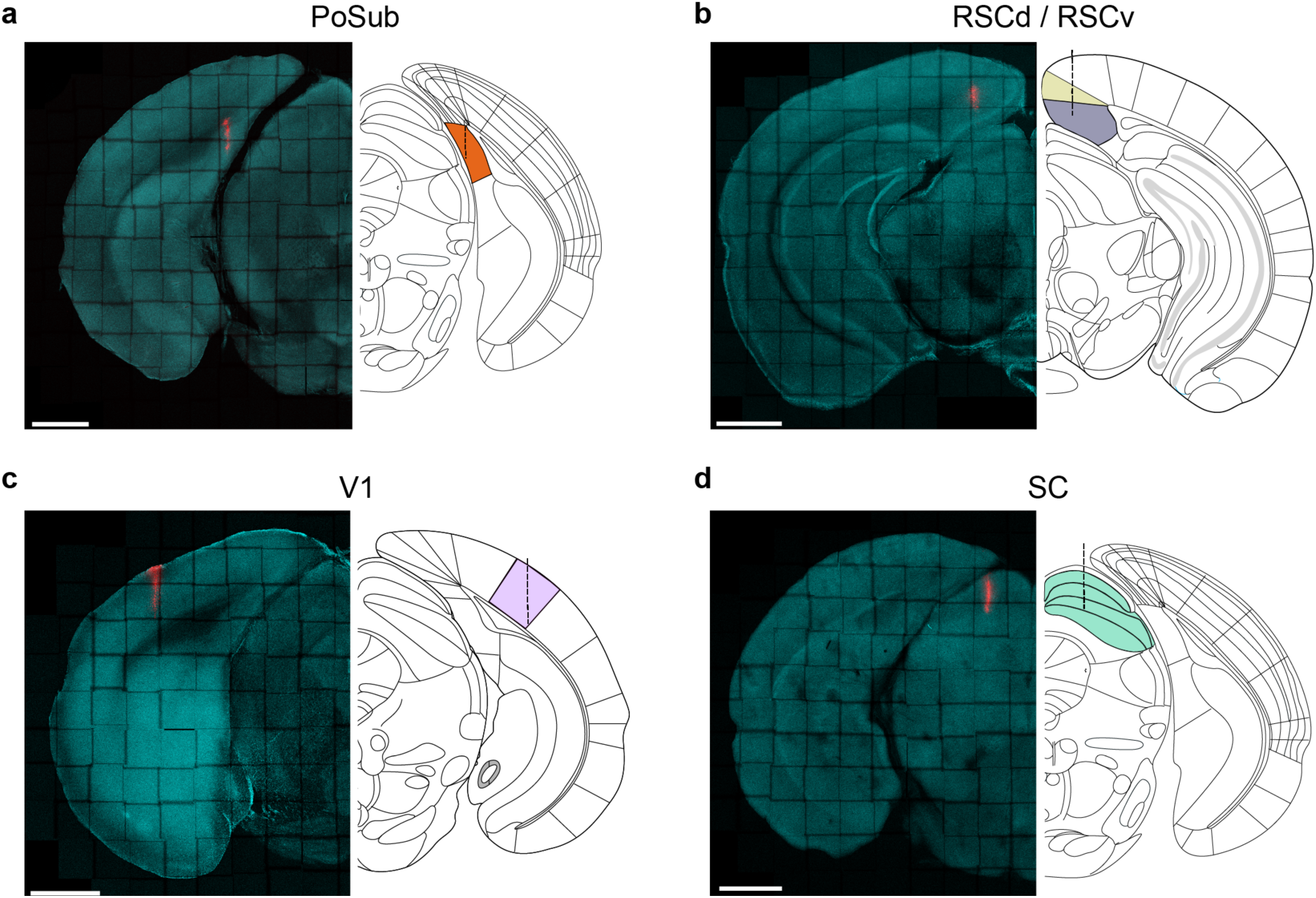
Coronal section showing insertion sites for electrophysiological experiments in Figure 2. **a-d**, Representative coronal sections showing the track of the electrophysiological probes dyed using a CM-DiI (red) for the five target regions. For reference, cell nuclei were stained using antibodies against DAPI (cyan). Scale bars represent 1 mm.

**Extended Data Fig. 6.**
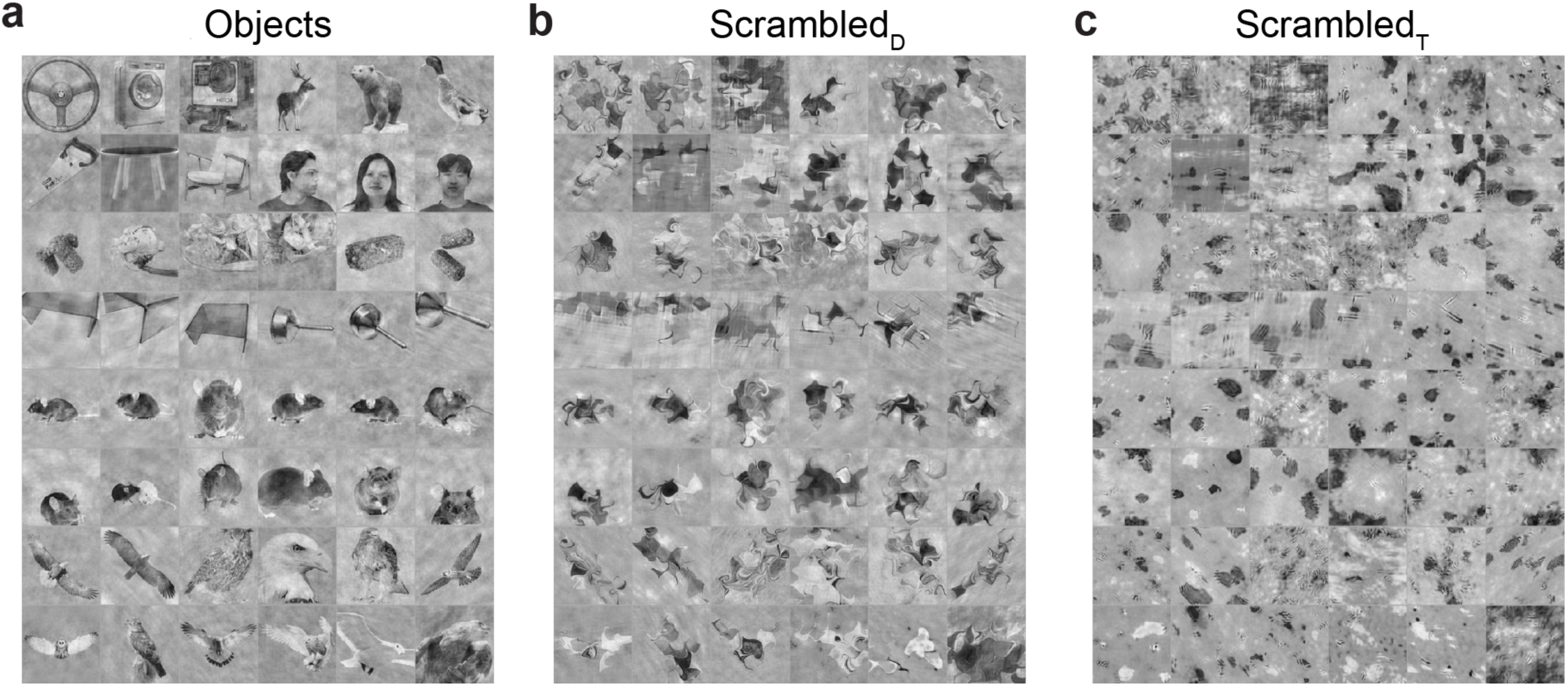
All objects and control stimuli used in electrophysiological experiments. **a**, The same 48 object images as in fUS experiments were presented in electrophysiological experiments. For electrophysiological experiments the images were presented individually (not in blocks as for fUS) and additional illuminance and spatial frequency matching was performed across all stimuli (see methods). **b**, The same 48 object images scrambled using the diffeomorphic transformation algorithm^38^. **c**, Again the same 48 objects images scrambled using a texture synthetizing algorithm^33^.

**Extended Data Fig. 7.**
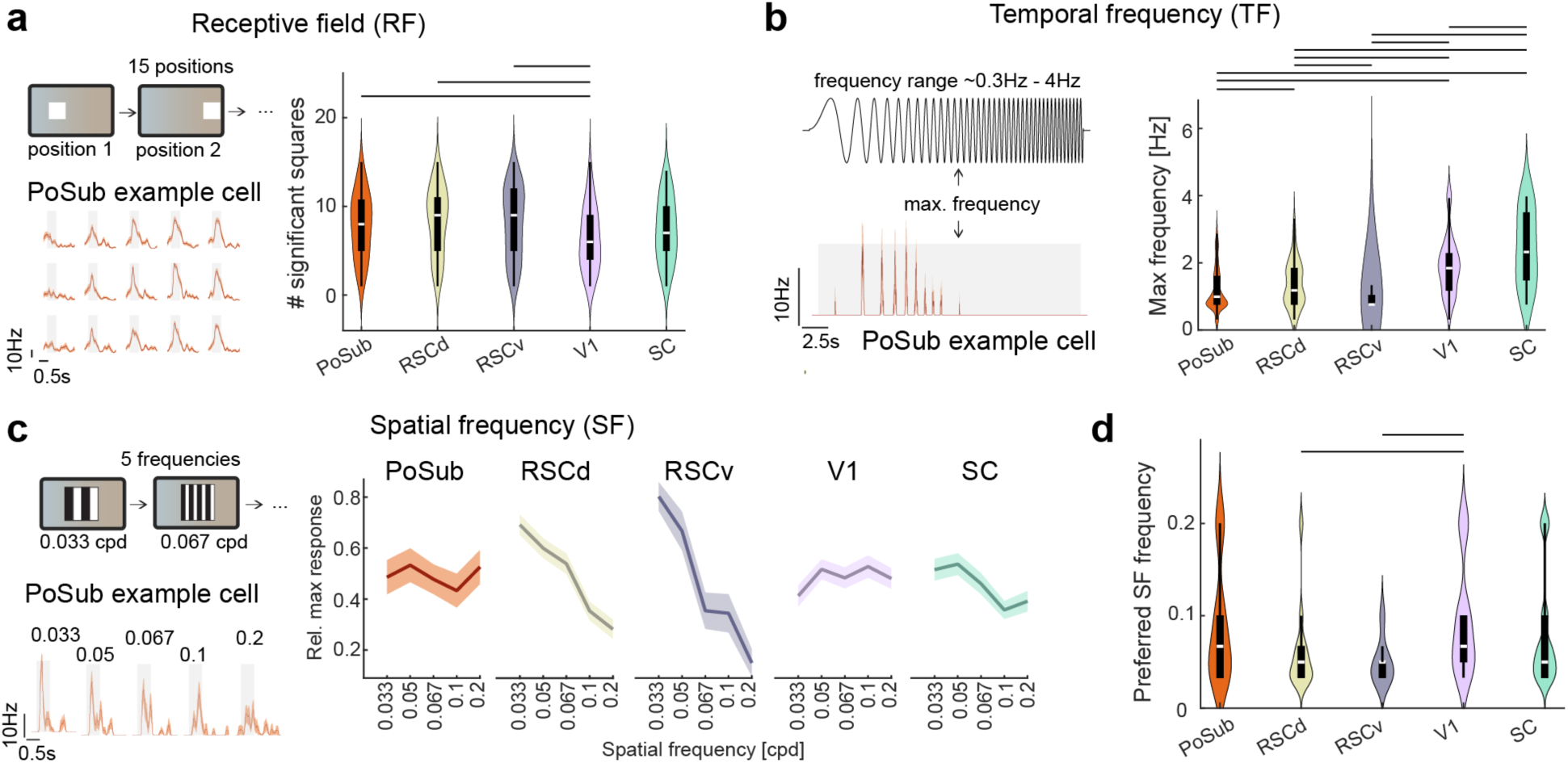
Neurons in spatial navigation areas have larger receptive fields and respond less well to high temporal frequencies. **a**, A white square was randomly presented at 15 different positions of the screen. *Bottom left*, Trial-averaged responses (mean ± s.e.m.) of a single cell in PoSub that responded to the majority of the 15 stimulus positions (gray boxes represent stimulus presentation). *Right*, Quantification of the number of squares that elicited a significant response for all analyzed cells. Significant differences are indicated by a black horizontal line, p < 0.05 (all p values are in Supplementary Table 2), B-H corrected, Mann-U-Whitney test, (n (from left to right) = 195, 165, 135, 210, 128 cells. **b**, Full-field chirp stimulus with increasing frequency was presented to the mouse, lasting 20 s. *Bottom left*, Trial-averaged response (mean ± s.e.m.) of an example cell in PoSub (gray box indicates duration of the chirp). *Right*, Quantification of the maximum frequency for all analyzed cells of indicated regions. Significant differences are indicated by a black line, p < 0.05 (all p values are in Supplementary Table 2), B-H corrected, Mann-Whitney U-test, (n (from left to right = 128, 114, 67, 167, 94). **c**, Different spatial frequencies (SFs) were presented at two orthogonal orientations (not shown). *Bottom left*, Trial-averaged responses (mean ± s.e.m.) to five SFs of a representative cell in PoSub indicating the relatively weak tuning and tendency to low SF preference observed in PoSub. *Right*, For each cell the responses were normalized to the maximal response and the mean ± s.e.m. across cells is displayed here (n (from left to right = 34, 81, 21, 89, 88 cells). **d**, Distributions of preferred SF frequency across all regions showing few significant differences indicated by a black line, p < 0.05 (all p values are in Supplementary Tables 2-4), B-H corrected, Mann-Whitney U-test (n’s as in c).

**Extended Data Fig. 8.**
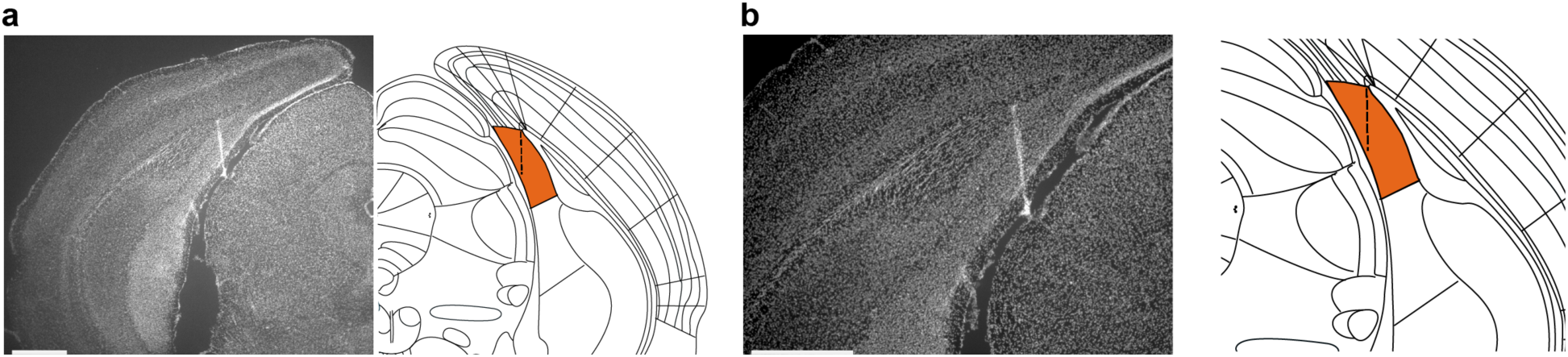
Coronal section showing position of chronically implanted silicon probe in PoSub. **a-b**, Overview (**a**) and zoom in (**b**) of an example coronal section stained with DAPI. The tract of the silicon probe is visible as a line of high intensity DAPI staining due to tissue scaring. Scale bars represent 1mm.

**Extended Data Fig. 9.**
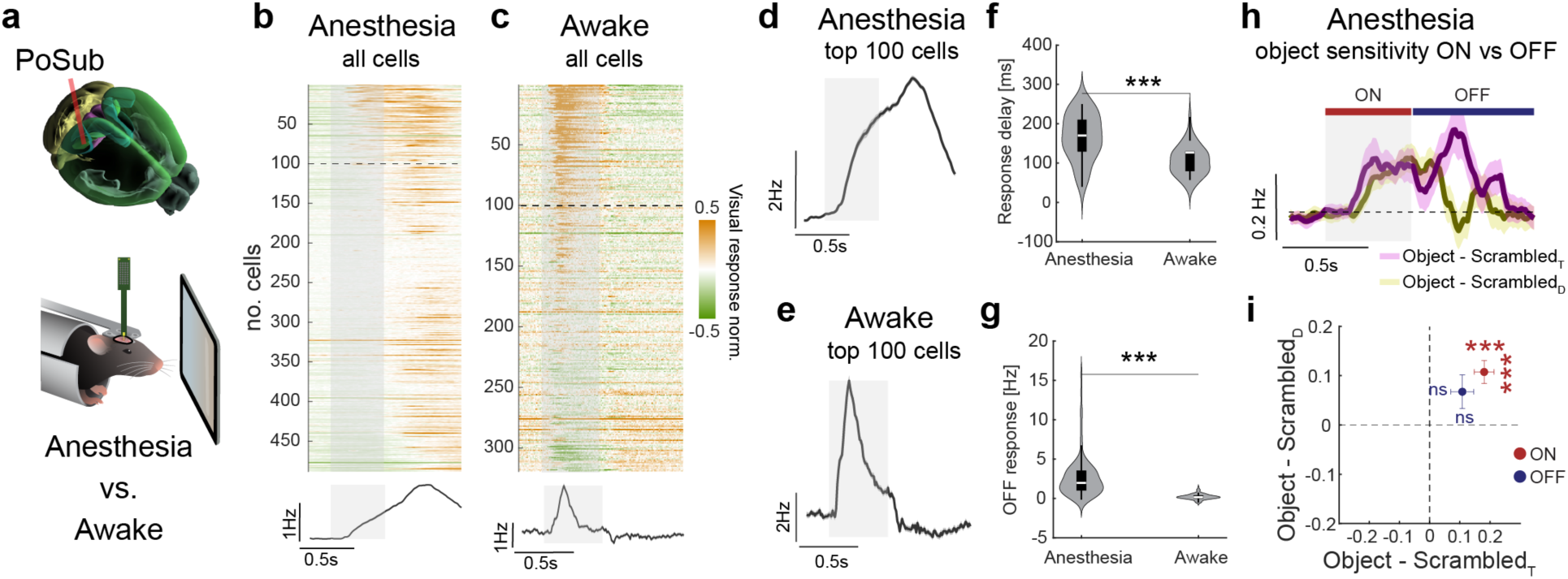
State-dependent differences of visual response properties in PoSub. **a**, Comparison of light-evoked electrophysiological recordings in PoSub from anesthetized animals (Fig. 2) and awake animals (Fig. 3/4). **b-c**, Shown are the normalized, trial-averaged responses (across all images) of all recorded cells in both conditions. Bottom panels show the average across all cells, showing slower and more sustained light-evoked responses under anesthesia. Under anesthesia, no negatively modulated cells were found and a large fraction of cells showed only responses after stimulus offset. **d-e**, To directly compare response dynamics between conditions, the top 100 responding cells (dashed line in b,c) were selected and the average of these cells is shown. **f**, Cells in PoSub respond significantly slower to visual stimuli under anesthesia than in awake condition, p = 1.5*10^-10^, Mann-Whitney U-test (n = 100/100 cells). **g**, In awake condition almost no OFF responses were observed, while cells had strong OFF responses under anesthesia, p=5.3*10^-25^, Mann-Whitney U-test (n=100/100 cells). Note that under anesthesia, the OFF response often merged with the slow ON response. **h,i**, As we found substantial OFF responses under anesthesia, we wondered whether the object sensitivity in PoSub described in Fig. 2 (significant difference of object responses in comparison to both scrambled image types) was restricted to the ON response. We found that in the OFF response, despite a trend towards a preference for Objects over ScrambledT, there was not a significant object preference. p (Object-ScrambledT, ON / OFF) = 9.6*10^-9^ / 0.097, p (Object-ScrambledD, ON / OFF) = 7.4*10^-7^ / 0.269, B-H corrected, Wilcoxon signed rank test (n = 192 cells).

**Extended Data Fig. 10.**
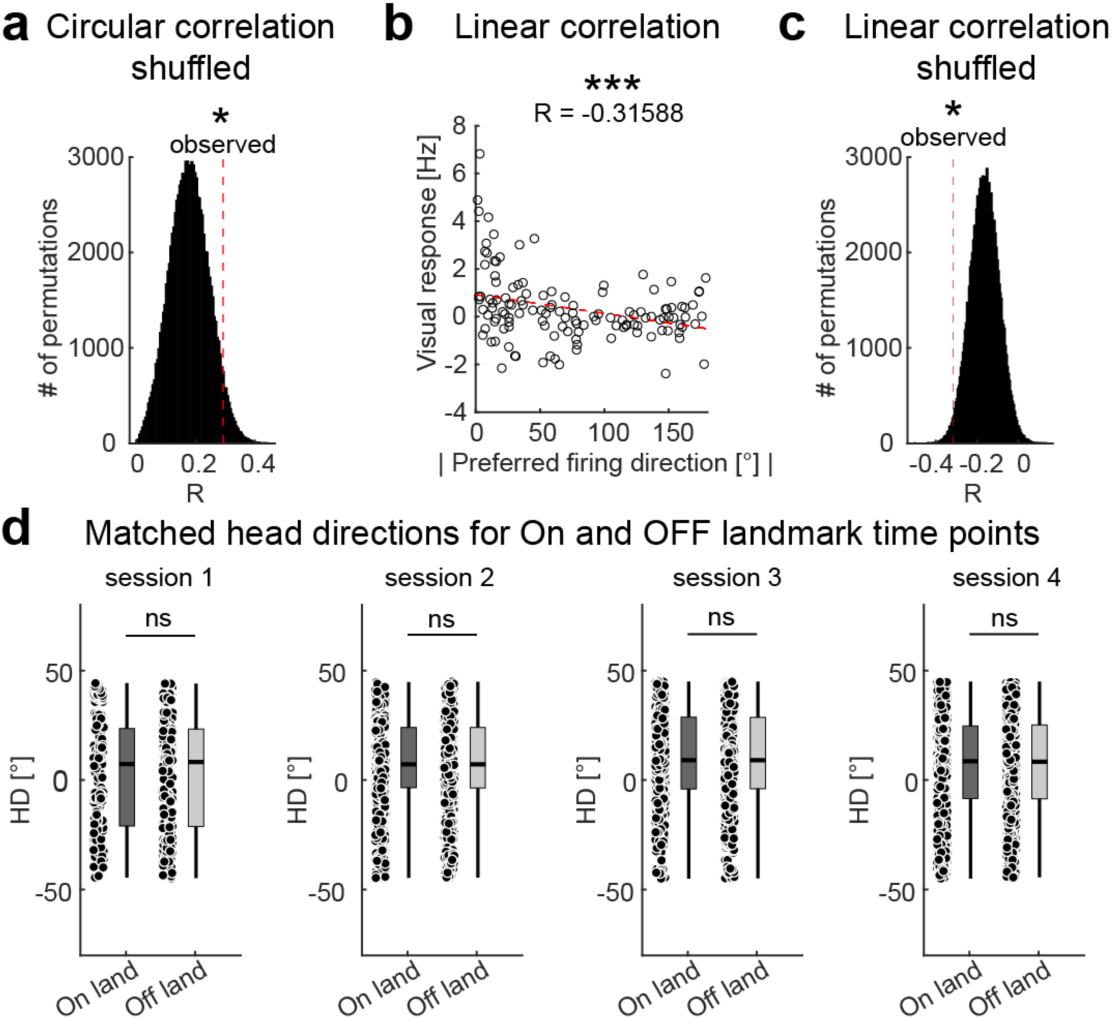
Analysis of HD population tuning and visual responses. **a**, Results of permutation test to ensure that the ring realignment procedure does not account for the circular correlation coefficients observed in Fig. 4. The histogram displays circular correlation coefficients after 100,000 permutations of visual response shuffling before ring alignment. The dotted red line indicates the observed circular correlation coefficient between visual response and preferred firing direction (p = 0.044). **b**, Pearson correlation between visual response and absolute preferred firing direction, p = 2.9*10^-4^ (n = 127 cells). Red dotted line represents a linear fit. **c**, Similar to (a), the distribution of Pearson correlation coefficients after shuffling the visual responses before ring alignments (100,000 permutations, p = 0.015). **d**, Related to Fig. 4f,g. To ensure that the differences we observed between the On and Off landmark activity were not driven by a potential bias in selection of time points (which could result in differences in firing activity due to head direction-dependent firing alone), we matched the head directions included in the analysis for each session, Mann-Whitney U-test (uncorrected), p (from left to right) = 0.988, 0.986, 0.99, 0.996 (n (from left to right) = 213/213, 751/ 751, 983/983, 1195/1195).

**Extended Data Fig. 11.**
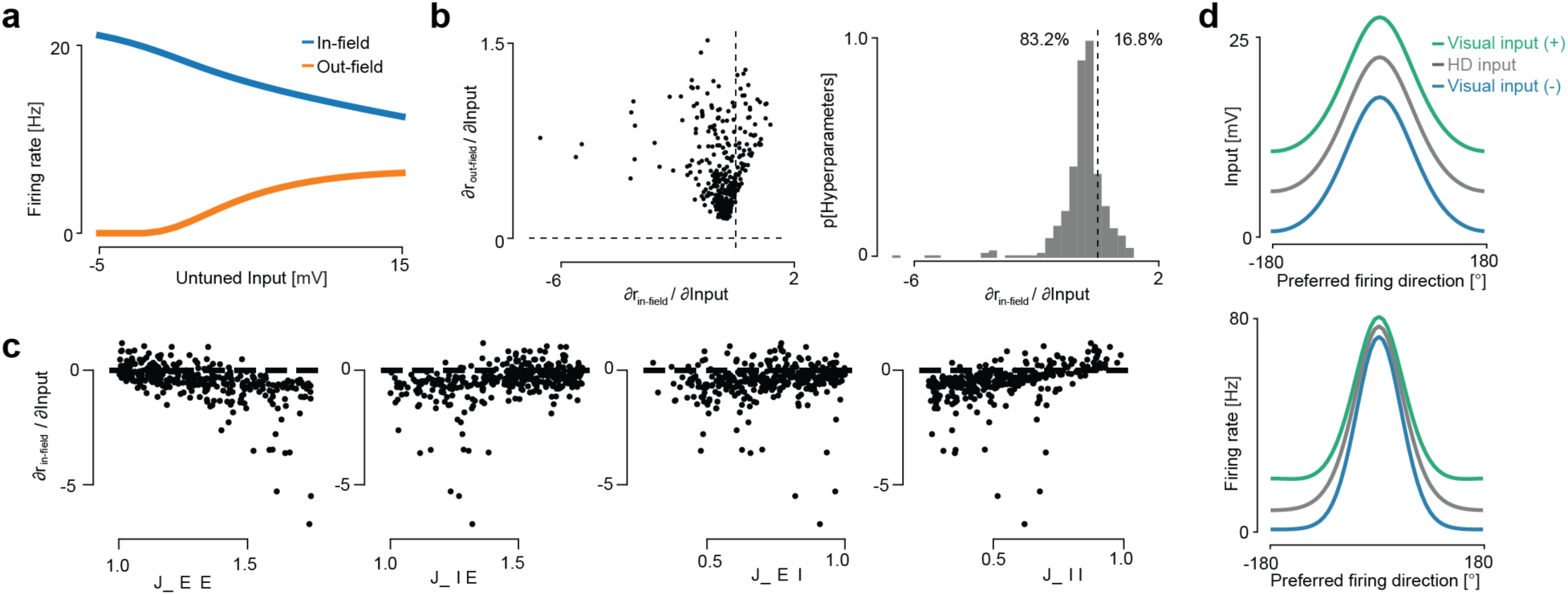
SSN model results are robust to hyperparameter variations. **a,** Equilibrium firing rate of in-field (model HD cells with a preferred firing direction of 0°, corresponding to the current HD) and out-field model units (corresponding to HD cells with a preferred firing direction of 180°) as a function of the level of untuned visual input. Increasing visual input always decreased the firing rate of in-field model units and increased the firing rate of out-field model HD neurons. Visual input only recapitulated our experimental finding if it resulted in a decrease in network input. **b**, The effect of varying the strength of untuned visual input on the firing rates of in-field (∂_rin-field_/∂Input) and out-field (∂_rout-field_/∂Input) model HD units over a population of networks with different randomly selected recurrent strength parameters. While the firing rate of out-field model HD units always increased as the visual input increased, the firing rate of in-field neurons decreased in most network instantiations as the visual input strength increased. **c**, ∂_rin-field/∂_Input for all network instantiations in the random parameter population, as a function of the E->E, E->I, I->E, and I->I weight. **d**, Same as Fig. 4l,m, but for inhibitory model units. Unlike the excitatory neurons, the firing rate of inhibitory model units always increased with increasing visual input.

**Supplementary Table 1.** All visually active regions and their network assignments based on literature review.

| Region name | Abbreviation | Mayor Division CCF | Assigned Network | Explanation/ References |
| --- | --- | --- | --- | --- |
| Anteromedial visual area | VISam | Isocortex | vision | higher visual area, adjacent to VISp <sup>69-72</sup> |
| Rostrolateral visual area | VISrl | Isocortex | vision | higher visual area, adjacent to VISp <sup>69-72</sup> |
| Anterior visual area | VISa | Isocortex | vision | higher visual area, adjacent to VISp <sup>69-72</sup> |
| Poteromedial visual area | VISpm | Isocortex | vision | higher visual area, adjacent to VISp <sup>69-72</sup> |
| Lateral visual area | VISl | Isocortex | vision | higher visual area, adjacent to VISp <sup>69-72</sup> |
| Posterolateral visual area | VISpl | Isocortex | vision | higher visual area, adjacent to VISp <sup>69-72</sup> |
| Anterolateral visual area | VISal | Isocortex | vision | higher visual area, adjacent to VISp <sup>69-72</sup> |
| Laterointermediate visual area | VISli | Isocortex | vision | higher visual area, adjacent to VISp <sup>69-72</sup> |
| Primary visual area | VISp | Isocortex | vision | primary visual cortical area, main target of LGd |
| Lateral group of the dorsal thalamus | LAT | Thalamus | vision | Two large nuclei of the LAT (LP and PO) receive direct input from the retina <sup>73,74</sup> |
| Dorsal part of the lateral geniculate complex | LGd | Thalamus | vision | Beside the SC, LGd is the main target of the retina <sup>75</sup> |
| Pretectal region | PRT | Midbrain | vision | Most nuclei of the PRT receive dense, direct input from the retina <sup>73</sup> |
| Peripeduncular nucleus | PP | Thalamus | vision | The PP receive dense, direct input from the retina <sup>73</sup> |
| Superior Colliculus | SC | Midbrain | vision | Superficial layers of the SC receive direct input from the retina <sup>75</sup> |
| brachium of the inferior colliculus | bic | fiber tracts | other | this fiber tract connects the inferior colliculus with the MG, two major aspect of the auditory system <sup>76</sup> |
| Infralimbic area | ILA | Isocortex | other | ILA is prefrontal region not particularly known for visual processing but for processing affective behaviors <sup>77,78</sup> |
| Medial geniculate complex | MG | Thalamus | other | Medial geniculate complex is a central aspect of the auditory system <sup>76</sup> |
| Periaqueductal gray | PAG | Midbrain | other | The PAG is diverse but is i.e. involved in defensive, social and emotional behaviors <sup>79-81</sup> |
| Ammon's horn, dorsal | CAd | Hippocampal formation | spatial navigation | The hippocampal formation is well known for its role in spatial navigation and memory <sup>82,83</sup> |
| Dentate gyrus, dorsal | DGd | Hippocampal formation | spatial navigation | The hippocampal formation is well known for its role in spatial navigation and memory <sup>82,83</sup> |
| Parasubiculum | PAR | Hippocampal formation | spatial navigation | The hippocampal formation is well known for its role in spatial navigation and memory <sup>82,83</sup> |
| Postsubiculum | POST | Hippocampal formation | spatial navigation | The hippocampal formation is well known for its role in spatial navigation and memory <sup>82,83</sup> |
| Area prostriata | APr | Hippocampal formation | spatial navigation | The hippocampal formation is well known for its role in spatial navigation and memory<sup>82,83</sup> |
| Subiculum | SUB | Hippocampal formation | spatial navigation | The hippocampal formation is well known for its role in spatial navigation and memory<sup>82,83</sup> |
| Restrosplenial area, dorsal | RSCd | Isocortex | spatial navigation | Similar to hippocampus, RSC is heavily studied in the context of spatial navigation<sup>6,84,85</sup> |
| Retrosplenial cortex, agranular area | RSCagl | Isocortex | spatial navigation | Similar to hippocampus, RSC is heavily studied in the context of spatial navigation<sup>6,84,85</sup> |
| Restrosplenial area, ventral | RSCv | Isocortex | spatial navigation | Similar to hippocampus, RSC is heavily studied in the context of spatial navigation<sup>6,84,85</sup> |

**Supplementary Table 2.** P values for statistical comparisons of RF sizes (Extended Data Fig. 7a)

|  | PoSub | RSCd | RSCv | V1 | SC |
| --- | --- | --- | --- | --- | --- |
| PoSub | | 0.72 | 0.841 | $3.1 \times 10^{-4}$ | 0.72 |
| RSCd | | | 0.841 | $5.9 \times 10^{-6}$ | 0.129 |
| RSCv | | | | $3.7 \times 10^{-4}$ | 0.37 |
| V1 |  |  |  |  | 0.129 |
| SC |  |  |  |  |  |

**Supplementary Table 3.** P values for statistical comparisons of maximal frequency (Extended Data Fig. 7b)

|  | PoSub | RSCd | RSCv | V1 | SC |
| --- | --- | --- | --- | --- | --- |
| PoSub | | **0.045** | 0.08 | $5.2 \times 10^{-12}$ | $8.5 \times 10^{-15}$ |
| RSCd | | | **0.002** | $1.2 \times 10^{-6}$ | $1.4 \times 10^{-10}$ |
| RSCv | | | | $1.8 \times 10^{-11}$ | $3.9 \times 10^{-13}$ |
| V1 |  |  |  |  | **0.002** |
| SC |  |  |  |  |  |

**Supplementary Table 4.** P values for statistical comparisons of preferred SF (Extended Data Fig. 7d)

|  | PoSub | RSCd | RSCv | V1 | SC |
| --- | --- | --- | --- | --- | --- |
| PoSub |  | 0.092 | 0.091 | 1.183 | 1.183 |
| RSCd | | | 0.795 | $1.9 \times 10^{-5}$ | 0.097 |
| RSCv |  |  |  | **0.002** | 0.105 |
| V1 |  |  |  |  | 0.092 |
| SC |  |  |  |  |  |

**Supplementary Table 5.** Default parameters used for the SSN model.

| Parameter | Value | Random Sweep Range |
| --- | --- | --- |
| $\tau_\theta$ | E: 20ms, I: 10ms | |
| $V_{rest}$ | -70 mV | |
| $\ell_{syn}$ | 45 degrees | |
| $W_{EE}$ | 1.25 | [1.00, 1.75] |
| $W_{IE}$ | 1.2 | [1.00, 1.75] |
| $W_{EI}$ | 0.65 | [0.25, 1.00] |
| $W_{II}$ | 0.5 | [0.25, 1.00] |
| $A_{max}$ | 20 Hz | |
| $\ell_{stim}$ | 60 degrees | |
| $n$ | 2.0 | |
| $V_0$ | -70 mV | |
| $k$ | 0.3 | |

